# Lactate treatment improves brain biochemistry and cognitive function in transgenic Alzheimer’s and wild-type mice

**DOI:** 10.64898/2026.01.28.702254

**Authors:** Imen Belhaj, Ingrid Åmellem, Can H. Tartanoglu, Hanne M. Weidemann, Evan M. Vallenari, Mingyi Yang, Farrukh A. Chaudhry, Magnar Bjørås, Jon Storm-Mathisen, Linda Hildegard Bergersen

## Abstract

Lactate, a well-known metabolite and signalling molecule, holds therapeutic potential for neurodegenerative diseases. Here, we investigated the effects of chronic lactate treatment on cognition and molecular biomarkers in the 5XFAD mouse model of Alzheimer’s disease (AD) and in wild-type (WT) controls using behavioural testing alongside proteomic and transcriptomic analyses. Mice received lactate or vehicle injections 4 days per week for 11 weeks, with behavioural testing before and after the treatment period. Lactate improved working memory in late-treated AD mice, without eliciting anxiety-like behaviour. At the molecular level, lactate reduced *Il1b* expression, and in a sex-dependent manner, normalised NEFL, and enhanced synaptic integrity proteins (OPCML, PPFIA2, STXBP3, SYT1, VGLUT2, VSNL1) in AD mice, while also augmenting mitochondrial regulators (ATP5G2, GRPEL1, SLC25A23) across genotypes. Notably, lactate upregulated low-abundance ionotropic glutamate receptor mRNAs (*Grik3, Grin2c, Grid2ip*) in female AD mice, indicating enhanced glutamatergic signalling. In WT mice, lactate increased expression of neurotrophic factors (*Bdnf, Igf1, Vegfa*), anti-inflammatory cytokines (*Il4* and *Il13*), and the neuronal lactate transporter *Mct2*, suggesting promoted neuronal resilience. Together, these findings indicate that lactate treatment can mitigate cognitive decline and enhance molecular pathways of resilience in AD, warranting larger, age-stratified studies to validate its therapeutic potential and elucidate underlying mechanisms.

## Introduction

Alzheimer’s disease (AD) is the most prevalent form of dementia, imposing a significant and increasing burden on individuals and society, particularly as life expectancy rises. Despite the growing prevalence of AD and other neurodegenerative dementias, there are currently limited established disease-modifying treatments or effective medications available^1^. In contrast, there is broad consensus that physical exercise exerts protective effects on the brain. Numerous studies in animal models and humans demonstrate that physical activity slows progression of AD and other neurodegenerative disorders^2–5^. A recent population-based prospective cohort study in Norway, known as the Trøndelag Health Study (HUNT study), has provided compelling evidence that increasing cardio-pulmonary fitness over time is associated with a substantial reduction in the risk of developing AD and in mortality rates^2,6^. These findings have sparked interest in lifestyle-based interventions, such as physical exercise, even when initiated later in life.

Our recent work^7^ has shed light on the role of lactate, acting largely through the hydroxycarboxylic acid receptor 1 (HCAR1), in bringing exercise-induced benefits to the brain including angiogenesis, neurogenesis and the upregulation of growth factors. The discovery suggests that exogenous lactate could potentially complement physical activity to maximise effects of exercise, particularly in older individuals and those with limited mobility^8,9^. Given the urgent need for new strategies to combat AD, understanding whether the lactate-mediated body-brain mechanism can serve as a viable approach is paramount. Although lactate has been identified as a key mediator of exercise-induced brain plasticity and neuroprotection through HCAR1 signalling, its direct and chronic effects in AD models remain unexplored. Addressing this gap is critical, as lactate could provide a mechanistic link between exercise and resilience to AD pathology.

Here, we hypothesised that chronic exogenous lactate administration would mitigate cognitive decline and modulate molecular markers of neurodegeneration and inflammation in the 5XFAD mouse model of AD. Because AD exhibits marked sex differences in prevalence and progression, sex was included as a biological variable in the study design. Since this is the first study to test lactate treatment in this AD model, we employed a broad exploratory approach combining behavioural, proteomic and transcriptomic analyses to capture potential effects across multiple biological levels.

## Results

### Lactate administration elevates blood lactate in mice

The AD mice used were hemizygous transgenic (Tg) mice expressing mutant amyloid precursor protein (*App)* and presenilin-1 (*Psen1)* (i.e. the 5XFAD model), along with their non-transgenic wild-type (WT) littermates. Treatment with lactate or vehicle was administered four times weekly to mice in the designated groups, over a duration of 11 weeks (Fig. 1a). Treatments commenced at two age intervals: early (at 1.1–2.3 months), and late (at 2.6–3.2 months) (Fig. 1b).

**Figure 1.**
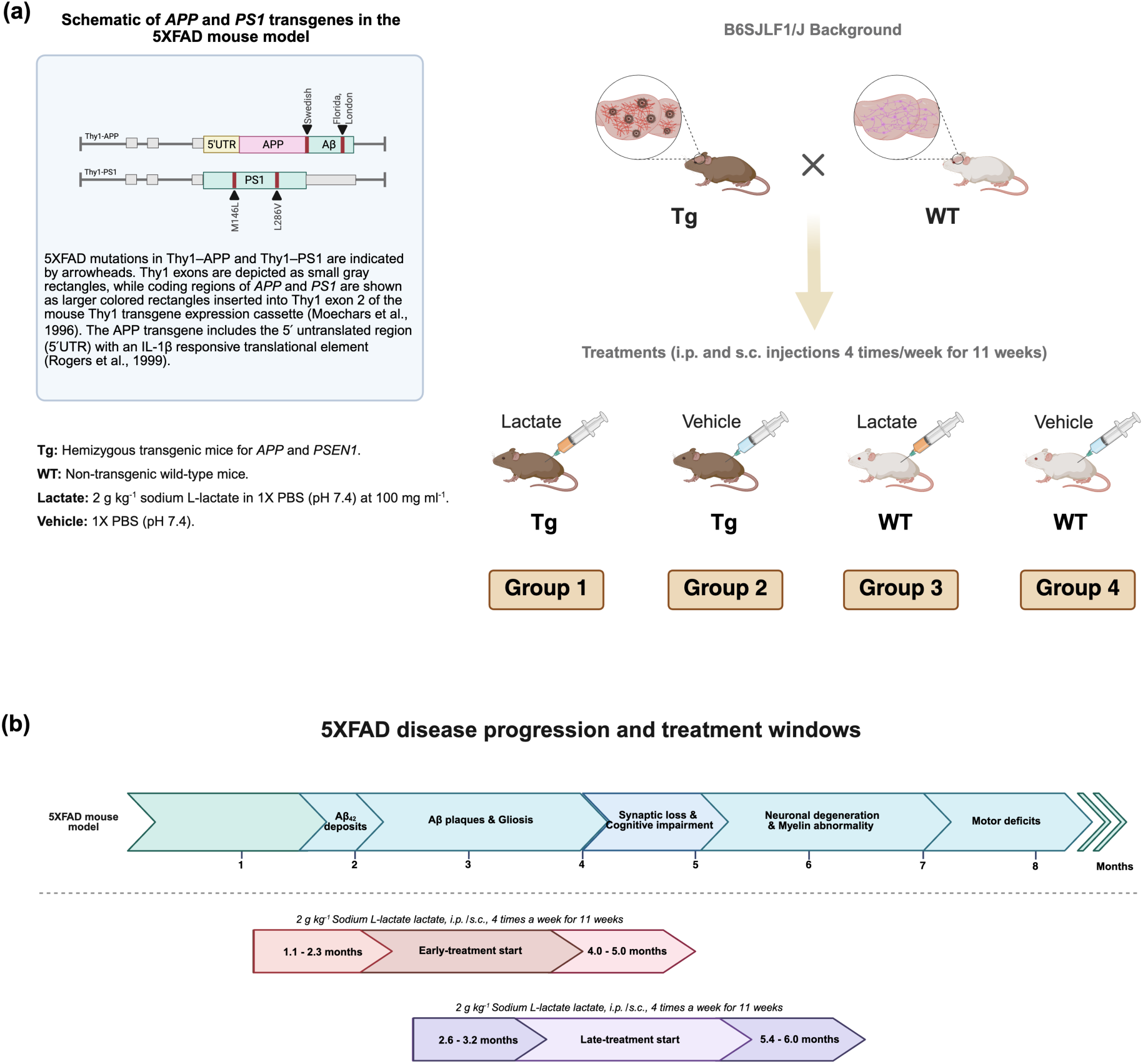
Experimental setup and disease progression in 5XFAD mice. **(a)** Schematic overview of the experimental setup and group allocation. Transgenic (Tg) 5XFAD mice, hemizygous for five familial Alzheimer’s disease (FAD) mutations on a B6JLF1/J background, and with wild-type (WT) littermates were divided into subgroups receiving lactate or vehicle. Treatments were administered via intraperitoneal (IP) and subcutaneous (SC) injections four times per week for 11 weeks, yielding four experimental groups. The schematic (upper left) illustrates insertion of amyloid precursor protein (*App*) and presenilin-1 (*Psen1*) transgenic mutations into exon 2 of the *Thy1* gene in the 5XFAD mouse model. **(b)** Timeline of AD-like progression in 5XFAD mice (in months), highlighting the two treatment windows: early- and late-treatment starts, corresponding to different disease stages at onset and completion. Figures created with BioRender.com.

Mice in the lactate-treated groups received 2 g kg⁻¹ sodium L-lactate at a concentration of 100 mg mL⁻¹ dissolved in 1× phosphate-buffered saline (PBS) (pH 7.4). To minimise local tissue reactions and stress while ensuring consistent systemic exposure, injection routes alternated between intraperitoneal (IP) and subcutaneous (SC). Our previous work has shown that these two routes yield comparable systemic lactate profiles in mice^10^, supporting their equivalence under our experimental conditions.

Blood lactate levels peaked to ∼23 mM at 13 min post-injection and declined to ∼15 mM at 37 min. For comparison with our previous work^7^, we also tested a higher concentration (200 mg mL⁻¹). This pilot test yielded blood lactate levels similar to those observed with the 100 mg mL⁻¹ dose (Supplementary Fig. 1), with peak levels likewise around 23 mM, but occurring earlier (at 5 min post-injection). To better characterise clearance dynamics, the decline phase (13–37 min) was fitted to a one-phase exponential decay model with the plateau constrained to 3 mM, corresponding to physiological baseline. This analysis yielded a rate constant k = 0.036 min⁻¹, a time constant (τ) of 27.7 min, and a half-life of ∼19 min. Based on this fit, lactate concentrations are projected to approach baseline after ∼3 half-lives (∼60 min), representing the practical return to baseline, while the asymptotic baseline is reached after ∼160 min. Thus, lactate elevation is transient, with effective normalisation within ∼1 h.

### Health monitoring of the mice

The body weights of the mice were monitored over the 12-week treatment period to assess their physical health status (Supplementary Fig. 2). As expected, body weight increased over time in all groups, with males exhibiting significantly higher weights than females (Supplementary Fig. 2a). Tg mice displayed slightly lower body weight than WT mice (Supplementary Fig. 2c). Similarly, lactate-treated mice had slightly lower body weight than vehicle-treated mice (Supplementary Fig. 2d). However, these differences were not statistically significant, suggesting that all experimental groups were in comparable physical condition.

### Lactate treatment protects against impaired cognitive performance

Working memory, assessed by spontaneous alternation in the Y-maze (Fig. 2), was significantly reduced in vehicle-treated AD mice (*p* = 0.0053 vs Tg lactate; *p* = 0.042 vs WT vehicle), reflecting progression of cognitive impairment, compared to AD mice treated with lactate. This protective effect of lactate was observed only in mice with late-treatment start (Fig. 2a), and not in those treated and tested during the early phase (Fig. 2b). Because no group differences emerged in the early-treatment cohort, testing likely preceded the onset of measurable cognitive impairment in AD mice^11,12^. When pre- and post-treatment performance was compared directly, late-treated Tg mice receiving vehicle showed a significant decline (*p* = 0.0018; Supplementary Fig. 3a). No significant pre–post changes were observed in Tg lactate, WT lactate, or WT vehicle groups, suggesting that lactate mitigated the decline otherwise occurring in Tg vehicle mice. In the early-treatment cohort, none of the groups showed significant pre-post change (Supplementary Fig. 3b).

**Figure 2.**
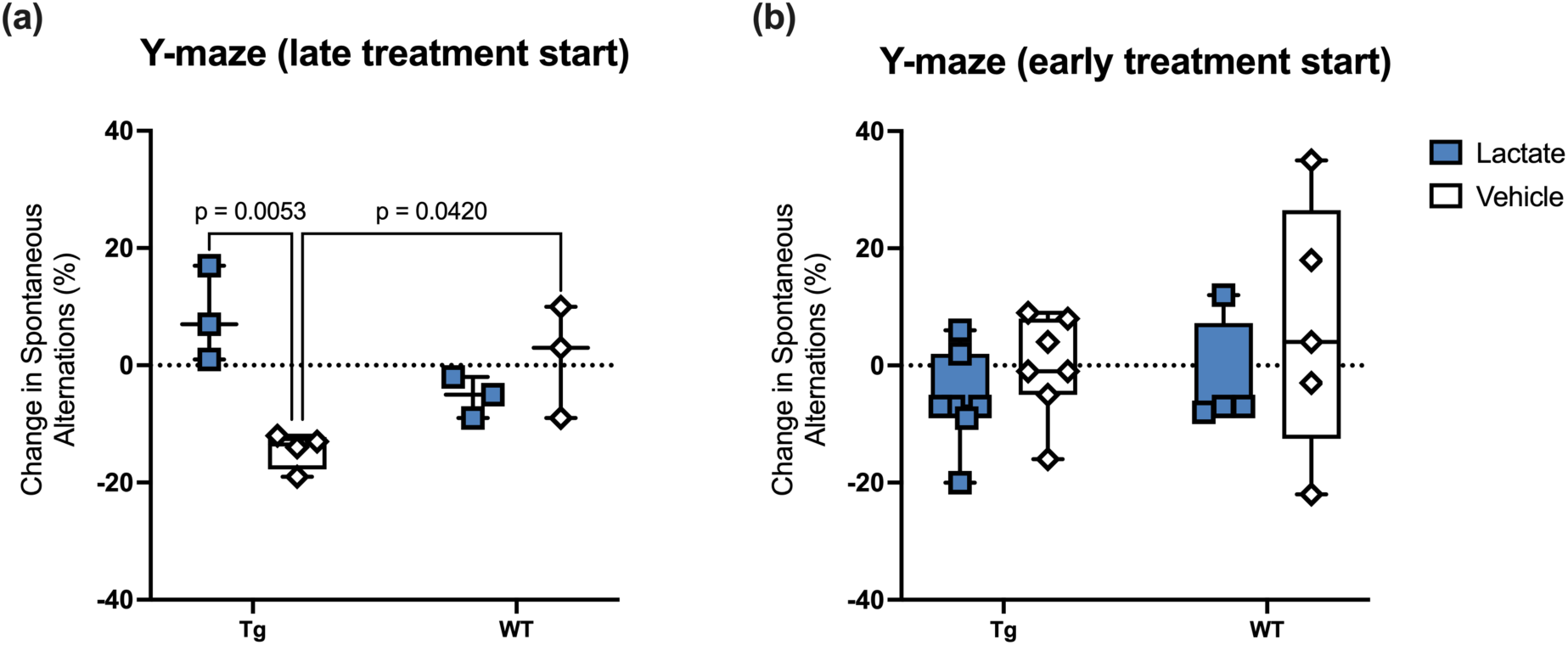
Lactate mitigates working memory decline in 5XFAD transgenic (Tg) mice in the Y-maze. The Y-maze was used to assess working memory before and after treatment of the same individual mouse, expressed as spontaneous alternation (SA) performance. Results are shown as change from baseline (%) for two treatment schedules, assessed in Tg and wild-type (WT) mice treated with lactate or vehicle. **(a)** Late-treatment start (2.6–3.2 months at the start; 5.4–6.0 months at the endpoint, both sexes): Two-way ANOVA revealed a significant genotype × treatment interaction. Tukey’s post hoc tests showed that Tg lactate-treated mice (*n* = 3) had higher SA than Tg vehicle controls (*n* = 4, *p* = 0.0053, *d* = 3.57). Tg vehicle mice also performed worse than WT vehicle mice (*n* = 3, *p* = 0.0420, *d* = 2.47). No differences were observed between lactate- (*n* = 3) and vehicle-treated WT groups. **(b)** Early-treatment start (1.1–2.3 months at the start; 4.0–5.0 months at the endpoint, both sexes): No significant treatment effects were detected in Tg mice (*n* = 7 per group) or WT mice (lactate: *n* = 4; vehicle: *n* = 5). Data are presented as box-and-whisker plots (median, interquartile range, all individual data points); dashed lines indicating baseline (0% change).

Notably, the number of spontaneous alternations in the Y-maze was not affected by hyperactive locomotion (Supplementary Fig. 4a–d) or by potential environmental factors drawing mice to specific maze regions (Supplementary Fig. 4e,f).

### No effect of lactate treatment on anxiety-like behaviour in the open-field test (OFT)

Lactate injections have been associated with acute anxiety attacks in susceptible human individuals^13^ and experimental animals^14^. In rodents, the interplay between their instinctive aversion to open areas, driven by perceived predator exposure, and their innate curiosity to explore new territories presents a unique opportunity to assess anxiety levels, locomotor function, and exploratory behaviour. To assess these parameters, we conducted OFT on all experimental groups. Our findings revealed no significant differences between groups (Supplementary Fig. 5), suggesting that lactate treatment, under our experimental conditions, did not induce anxiety-like behaviour and thus did not confound the behavioural outcomes.

Having established that the behavioural effects of lactate were not attributable to anxiety or locomotor confounds, we next investigated molecular correlates by performing proteomic profiling. This unbiased approach enabled assessment of broad protein-level changes induced by lactate treatment, with particular focus on synaptic and modulatory proteins relevant to cognition.

### Global proteomic analyses identify differentially expressed proteins related to AD, neuronal, mitochondrial, and metabolic processes

Given evidence for sex differences in AD pathology and treatment response, male and female mice were analysed separately. Whole brain hemisphere (hemibrain) liquid chromatography-tandem mass spectrometry (LC-MS/MS) profiling identified 8 differentially expressed proteins (DEPs) in females, and 8 in males (Fig. 3, detailed panels in Supplementary Fig. 7). Volcano plots for each comparison are shown in Fig. 3d–k, and the full list of DEPs is provided in Supplementary Data (All_DEPs_MS).

**Figure 3.**
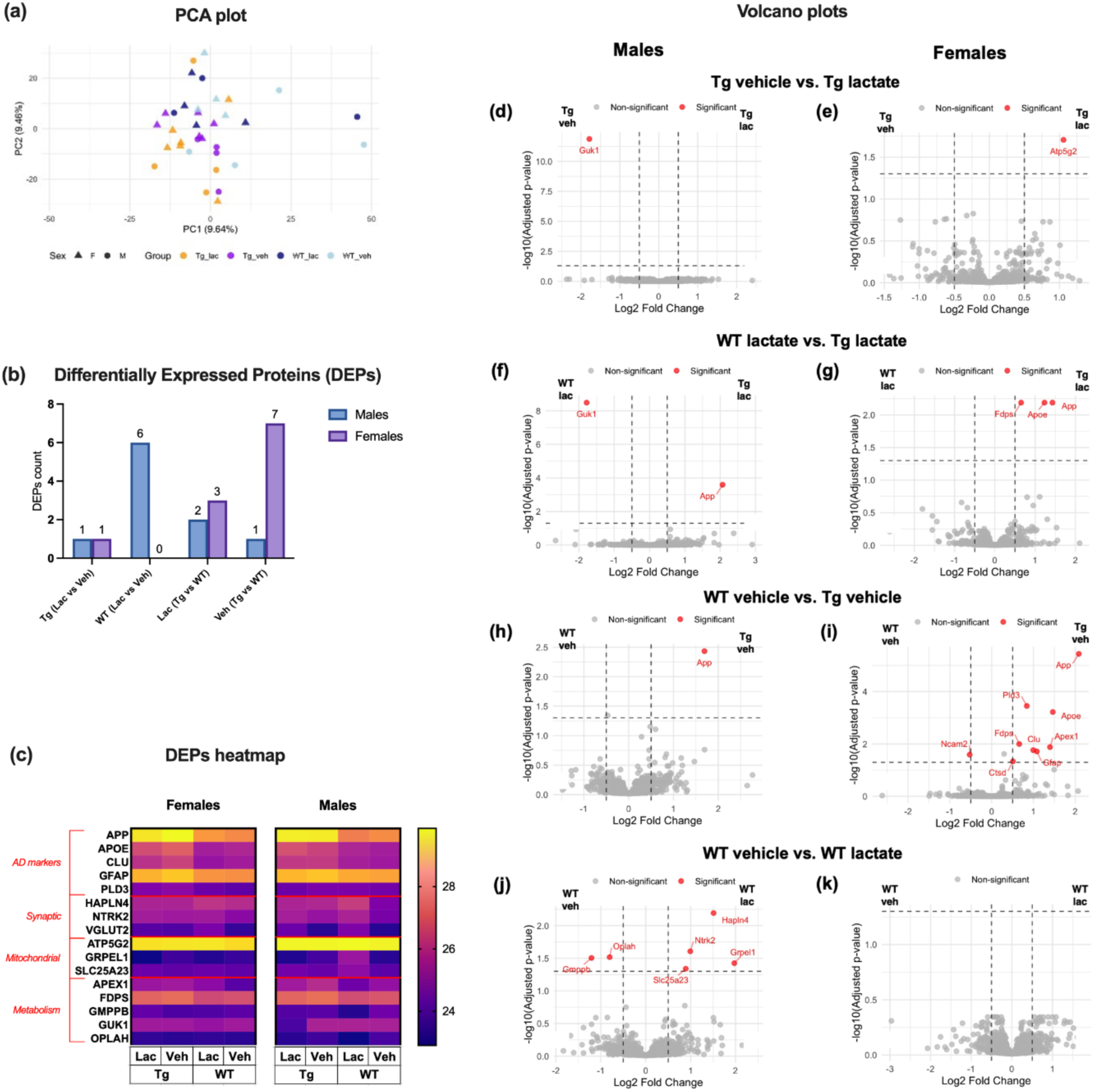
Global proteomic analyses and differentially expressed proteins (DEPs). Analyses were conducted in 5XFAD transgenic (Tg) and wild-type (WT) mice treated with lactate or vehicle. **(a)** Principal component analysis (PCA) of proteomic profiles, showing sample distribution along the first two principal components (PC1 and PC2), based on variance across 2,045 commonly detected proteins (unique genes). Shapes denote sex; colours denote genotype and treatment. **(b)** Bar plot showing the number of DEPs identified in lactate-treated versus vehicle-treated mice and in Tg versus WT, stratified by sex (blue = males; purple = females). DEPs were defined as proteins with adjusted *p* < 0.05 and absolute log₂ fold change > 0.5. **(c)** Heatmaps of DEPs grouped into four categories: AD markers (APP, APOE, CLU, GFAP, PLD3), synaptic proteins (HAPLN4, NTRK2, VGLUT2), mitochondrial proteins (ATP5G2, GRPEL1, SLC25A23), and metabolic regulators (APEX1, FDPS, GMPPB, GUK1, OPLAH). **(d–k)** Volcano plots of DEPs for pairwise comparisons: **(d)** male Tg vehicle vs Tg lactate, **(e)** female Tg vehicle vs Tg lactate, **(f)** male WT lactate vs Tg lactate, **(g)** female WT lactate vs Tg lactate, **(h)** male WT vehicle vs Tg vehicle, **(i)** female WT vehicle vs Tg vehicle, **(j)** male WT vehicle vs WT lactate, **(k)** female WT vehicle vs WT lactate. Significant DEPs are highlighted in red; dashed line indicates adjusted *p* = 0.05. Statistical analysis was performed using Wald statistics with Benjamini–Hochberg correction for multiple comparisons. The full protein list is provided in Supplementary Data.

Analysis revealed several key proteins related to AD pathology. Amyloid precursor protein (APP) and apolipoprotein E (APOE) were markedly increased in Tg versus WT mice, with no significant change with lactate treatment. Clusterin (CLU), a cytoprotective protein associated with AD and protein aggregation and glial fibrillary acidic protein (GFAP), an astroglial marker, were significantly elevated in Tg versus WT mice. Lactate did not significantly alter CLU or GFAP, although levels in lactate-treated Tg mice trended toward WT. Phospholipase D family member 3 (PLD3), an AD-risk gene product implicated in lysosomal function and amyloid-β (Aβ), was increased in female Tg relative to WT and unchanged by lactate (Fig. 3c; Supplementary Fig. 7a–e).

Lactate also increased hyaluronan and proteoglycan link protein 4 (HAPLN4) in WT males (*p* = 0.022) and neurotrophic receptor tyrosine kinase 2 (NTRK2) in WT females (*p* = 0.009) compared to vehicle controls (Fig. 3c; Supplementary Fig. 7f,g).

Mitochondrial proteins formed a notable subset of DEPs, being directly linked to neuronal energy metabolism and calcium buffering—both strongly implicated in AD. In Tg females, lactate increased ATP synthase membrane subunit c locus 2 (ATP5G2) (*p* = 0.026), a subunit of mitochondrial ATP synthase involved in neural energy production. In males, the mitochondrial protein transporter GRPEL1 (*p* = 0.009) and the mitochondrial Ca²⁺-transporter SLC25A23 (*p* = 0.056) were upregulated by lactate in WT but not Tg mice (Fig. 3c; Supplementary Fig. 7h–j).

Metabolic proteins were also differentially expressed. In WT females, lactate increased apurinic/apyrimidinic endodeoxyribonuclease 1 (APEX1) (*p* = 0.017). Farnesyl diphosphate synthase (FDPS) showed no significant effects of lactate, but a slight trend of reduction. In WT males, lactate reduced GDP-mannose pyrophosphorylase B (GMPPB) (*p* = 0.026) and 5-oxoprolinase / ATP-hydrolysing (OPLAH) (*p* = 0.006), while in Tg males it decreased guanylate kinase 1 (GUK1) (*p* = 0.002; Fig. 3c; Supplementary Fig. 7k–o).

### Lactate alters AD-related synaptic vesicle and structural protein expression in a sex-dependent manner

The proteins highlighted in Fig. 4 were selected for their established relevance to AD pathology, spanning neuronal integrity, synaptic function, and neurodegeneration.

**Figure 4.**
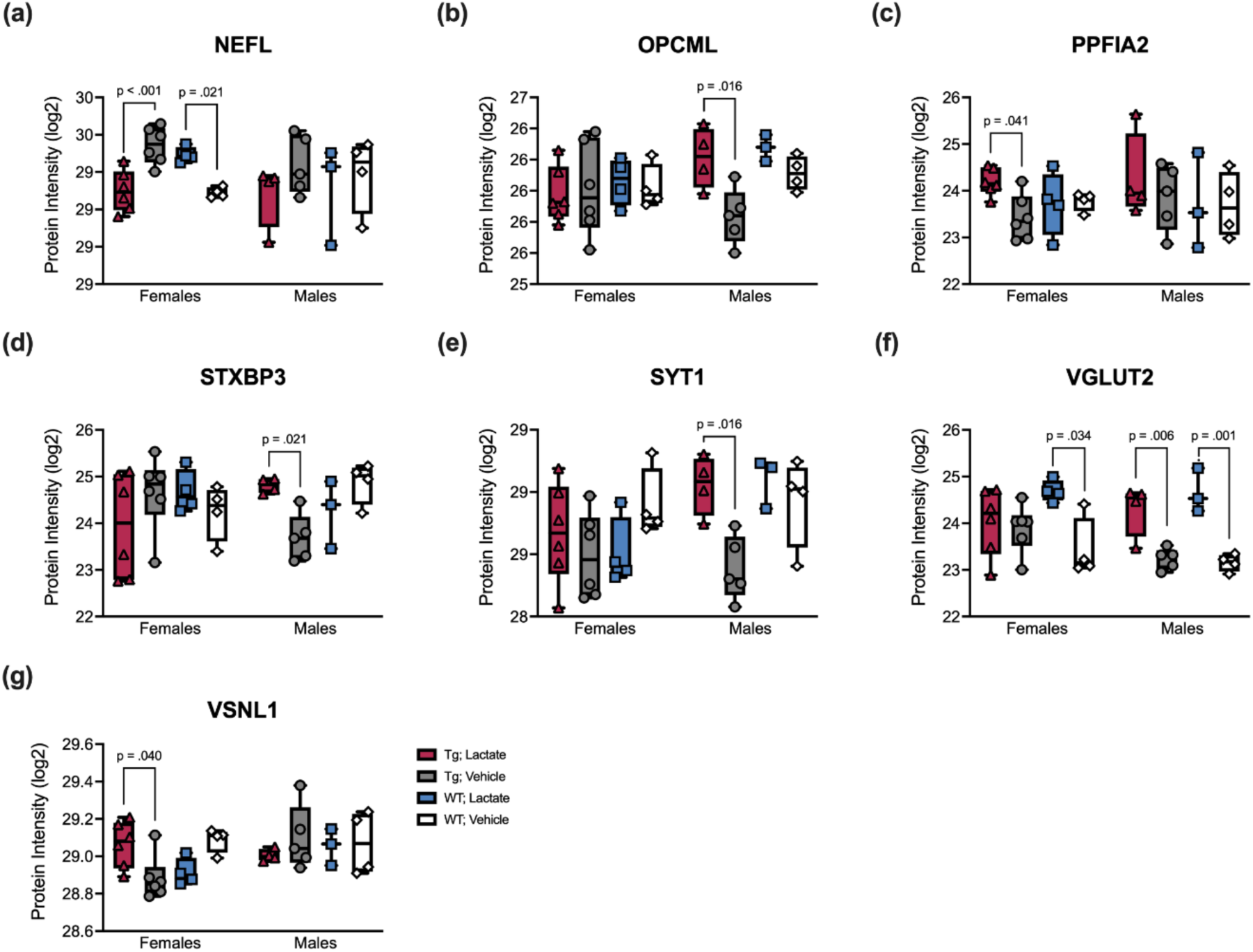
Lactate treatment modulates AD-related synaptic vesicle and structural proteins in a sex-dependent manner. Liquid chromatography-tandem mass spectrometry (LC-MS/MS) proteomics revealed changes in proteins central to presynaptic glutamate handling, vesicle release, calcium signalling, synaptic scaffolding, and axonal integrity in male and female mice, with early and late treatment groups combined, assessed in 5XFAD transgenic (Tg) and wild-type (WT) mice treated with lactate or vehicle. The experimental groups consisted of Tg mice receiving lactate (*n* = 10, 6 females and 4 males), Tg mice receiving vehicle (*n* = 11, 6 females and 5 males), WT mice receiving lactate (*n* = 7, 4 females and 3 males), and WT mice receiving vehicle (*n* = 8, 4 females and 4 males). Data are presented as box-and-whisker plots (median, interquartile range, and all individual data points). Statistical analyses were performed using one-way ANOVA followed by Tukey’s post hoc test; significant differences are indicated above the plots. **(a)** Neurofilament light polypeptide (NEFL), a cytoskeletal marker elevated in AD biofluids and tissue, was increased in WT females (*p* = 0.021, *d* = 2.31) and decreased in Tg mice treated with lactate (*p* < 0.001, *d* = 2.78), the latter suggesting a normalising effect of lactate on axonal integrity. **(b)** Opioid-binding protein/cell adhesion molecule-like (OPCML), a synaptic adhesion molecule, often reduced in AD, was upregulated in Tg males treated with lactate (*p* = 0.016, *d* = 2.42), consistent with improved synaptic adhesion and organisation. **(c)** Liprin-α2 (PPFIA2), a presynaptic active-zone scaffold that organises neurotransmitter release and is generally diminished in AD, was upregulated in Tg females with lactate treatment (*p* = 0.041, *d* = 1.71), consistent with strengthened presynaptic architecture. **(d)** Syntaxin-binding protein 3 (STXBP3), a regulator of SNARE-mediated vesicle release, typically reduced in AD, was upregulated in Tg males with lactate (*p* = 0.021, *d* = 2.33), pointing to enhanced vesicle release capacity. **(e)** Synaptotagmin-1 (SYT1), the Ca²⁺ sensor for vesicle fusion, frequently decreased in AD synapses, was increased in Tg males with lactate (*p* = 0.016, *d* = 2.45), suggesting restoration of presynaptic release capacity. **(f)** Vesicular glutamate transporter 2 (VGLUT2, also known as SLC17A6), required for loading glutamate into synaptic vesicles, typically reduced in AD, was upregulated in WT females (*p* = 0.034, *d* = 2.16), WT males (*p* = 0.001, *d* = 4.02), and Tg males (*p* = 0.006, *d* = 2.82), indicating enhanced presynaptic glutamate handling. **(g)** Visinin-like protein 1 (VSNL1), a neuronal Ca²⁺ sensor linked to AD pathology and neuronal signalling, was increased in Tg females with lactate (*p* = 0.040, *d* = 1.72), indicating lactate-linked modulation of Ca²⁺ signalling.

Neurofilament light polypeptide (NEFL), a marker of axonal damage, was elevated in Tg females receiving vehicle, but normalised with lactate (*p* < 0.001 vs Tg vehicle), whereas WT females treated with lactate showed increased NEFL compared to their WT controls (*p* = 0.021; Fig. 4a). Opioid-binding protein/cell adhesion molecule like (OPCML), typically reduced in AD, was upregulated in lactate-treated versus vehicle-treated Tg males (*p* = 0.016), suggesting enhanced synaptic adhesion and organisation (Fig. 4b). Liprin-α2 (PPFIA2), a presynaptic scaffold that organises neurotransmitter release, and reported reduced in AD, was increased in Tg females treated with lactate (*p* = 0.041), indicating strengthened presynaptic architecture (Fig. 4c). Syntaxin-binding protein 3 (STXBP3), involved in neurotransmitter release was upregulated in Tg males (*p* = 0.021; Fig. 4d). Synaptotagmin-1 (SYT1), a key Ca^2+^ sensor for synaptic vesicle fusion was likewise increased by lactate from low levels with vehicle in Tg males (*p* = 0.016; Fig. 4e). Vesicular glutamate transporter 2 (VGLUT2, also known as SLC17A6), a key mediator of excitatory neurotransmission, essential for brain development^15,16^, and downregulated in AD^17^, was increased in WT females (*p* = 0.034), WT males (*p* = 0.001) and Tg males (*p* = 0.006; Fig. 3c; 4f). Finally, Visinin-like protein 1 (VSNL1), a neuronal Ca^2+^ sensor protein linked to AD, was increased by lactate in Tg females (*p* = 0.040; Fig. 4g).

### Sex-specific differential gene expression patterns in response to lactate treatment

To investigate how lactate treatment influences brain transcriptional profiles, we performed RNA sequencing (RNA-seq) of hemibrains. We identified 230 differentially expressed genes (DEGs) in females and 145 in males (Fig. 5a, full list in Supplementary Data, All_DEGs_RNAseq). Heatmaps in Fig. 5b–e display prioritised lactate-regulated genes selected by largest fold-changes, multiple-testing significance, and biological relevance to AD pathways (glutamatergic signalling, inflammation, and autophagy), shown separately for Tg and WT within each sex.

**Figure 5.**
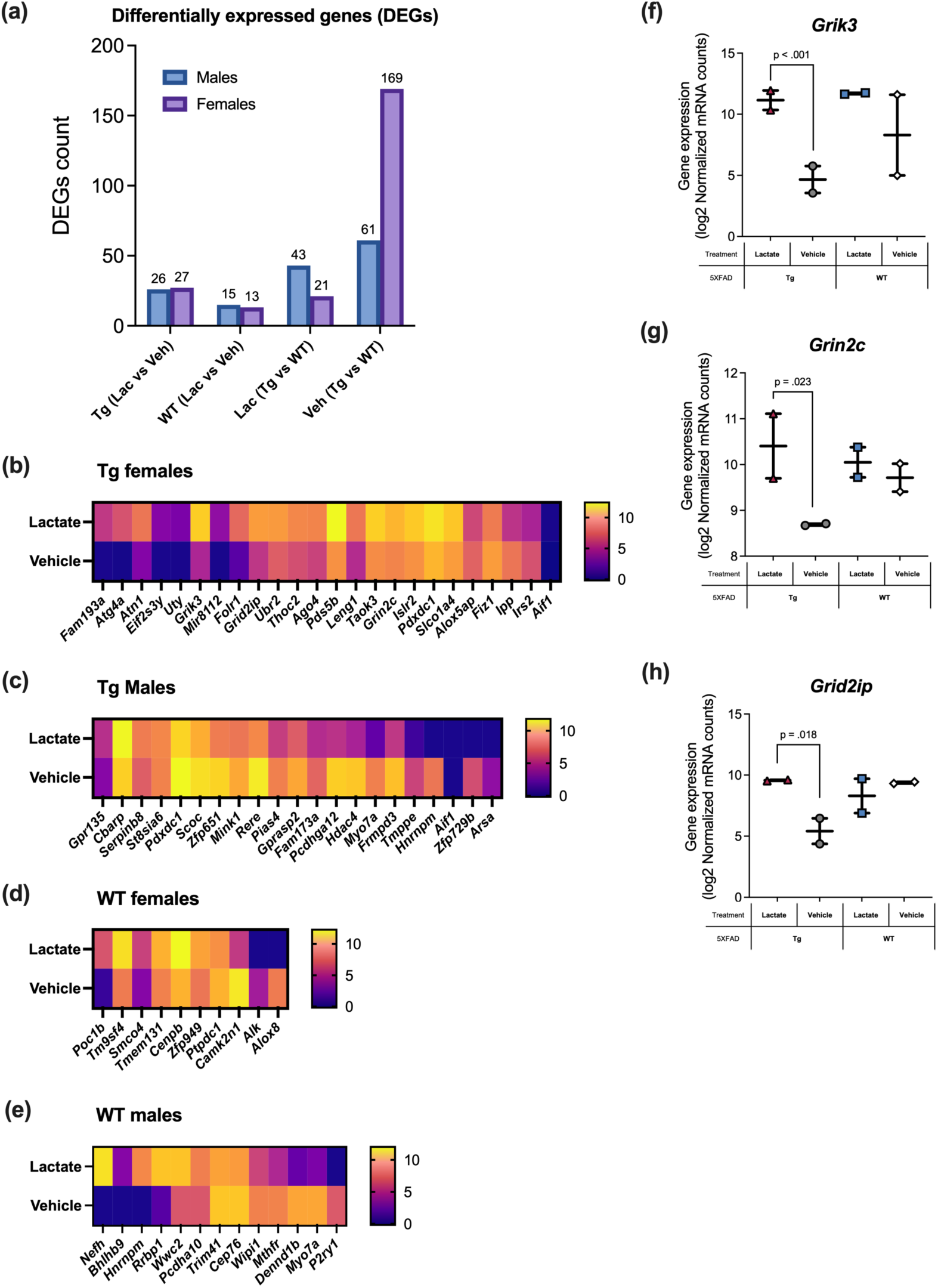
Sex-specific differential gene expression in response to early lactate treatment. RNA-seq analysis of 5XFAD transgenic (Tg) and wild-type (WT) mice treated with lactate or vehicle (early treatment: 1.1–2.3 months at the start, 4.0–5.0 months at the endpoint). **(a)** Bar plot showing the number of differentially expressed genes (DEGs) identified in lactate-versus vehicle-treated mice and in Tg versus WT, stratified by sex (blue = males; purple = females). DEGs were defined as transcripts with adjusted *p* < 0.05 and absolute log₂ fold change ≥ 1. **(b–e)** Heatmaps of log₂-transformed normalised expression values for DEGs (blue = lowest, yellow = highest), with genes ranked by log₂ fold change. Each group included *n* = 2 biological replicates, except WT male lactate (*n* = 3). All presented genes had adjusted *p* < 0.005. **(b)** DEGs regulated by lactate in female Tg mice. **(c)** DEGs regulated by lactate in male Tg mice. **(d)** DEGs regulated by lactate in female WT mice. **(e)** DEGs regulated by lactate in male WT mice. **(f–h)** Expression of selected glutamatergic genes significantly upregulated in lactate-treated Tg females. Data are presented as box-and-whisker plots (median, interquartile range, and all individual data points). **(f)** *Grik3* (*p* < 0.001), **(g)** *Grin2c* (*p* = 0.023), and **(h)** *Grid2ip* (*p* = 0.018). Statistical analysis was performed using Wald statistics with Benjamini–Hochberg correction for multiple comparisons. The full gene list is provided in Supplementary Data.

Transcriptomic analysis in female Tg mice revealed increased expression of three ionotropic glutamate receptor-related genes: *Grik3* (kainate receptor subunit; also known as *Eaa5*, *Glr7*, *Glur7*, *Gluk3*, *Glur7a*) (*p* < 0.001), *Grin2c* (NMDA receptor subunit; also known as *Nr2c*, *Glun2c*, *Nmdar2c*) (*p* = 0.023), and *Grid2ip* (glutamate receptor delta 2 interacting protein) (*p* = 0.018; Fig. 5f–h). Gene Ontology (GO) analysis of lactate-versus vehicle-treated female Tg mice showed significant enrichment driven by *Grik3* and *Grin2c* (Supplementary Fig. 6a,b). In male Tg mice, lactate-regulated DEGs were enriched in pathways related to epigenetic and post-translational regulation of transcription (Supplementary Fig. 6c,d). In WT mice, RNA-seq revealed no significant DEGs, likely due to small group sizes. However, real-time quantitative PCR (RT-qPCR) validation confirmed that late lactate treatment significantly increased *Grik3* (*p* = 0.021) and *Grin2c* (*p* = 0.015) mRNA in female WT mice (Supplementary Fig. 8a,b).

### Early administration of lactate ameliorates inflammation-associated gene expression

Motivated by proteomic pathways related to inflammation and synaptic modulation, we used targeted RT-qPCR in the early-treated cohort to examine candidate neuroinflammatory genes. Neuroinflammation is increasingly recognised as a driver of pathological progression in AD. As expected, the pro-inflammatory cytokine *Il1b* mRNA was elevated in Tg mice compared to WT and was normalised by lactate in Tg (*p* = 0.026; Fig. 6a). In WT, lactate treatment also increased the expression of the primarily anti-inflammatory cytokines *Il4* (*p* = 0.036; Fig. 6b) and *Il13* (*p* = 0.008; Fig. 6c). When early- and late-treatment cohorts were combined, results showed similar trends, but only *Il13* remained significant (*p* < 0.001; Supplementary Fig. 9a–c).

**Figure 6.**
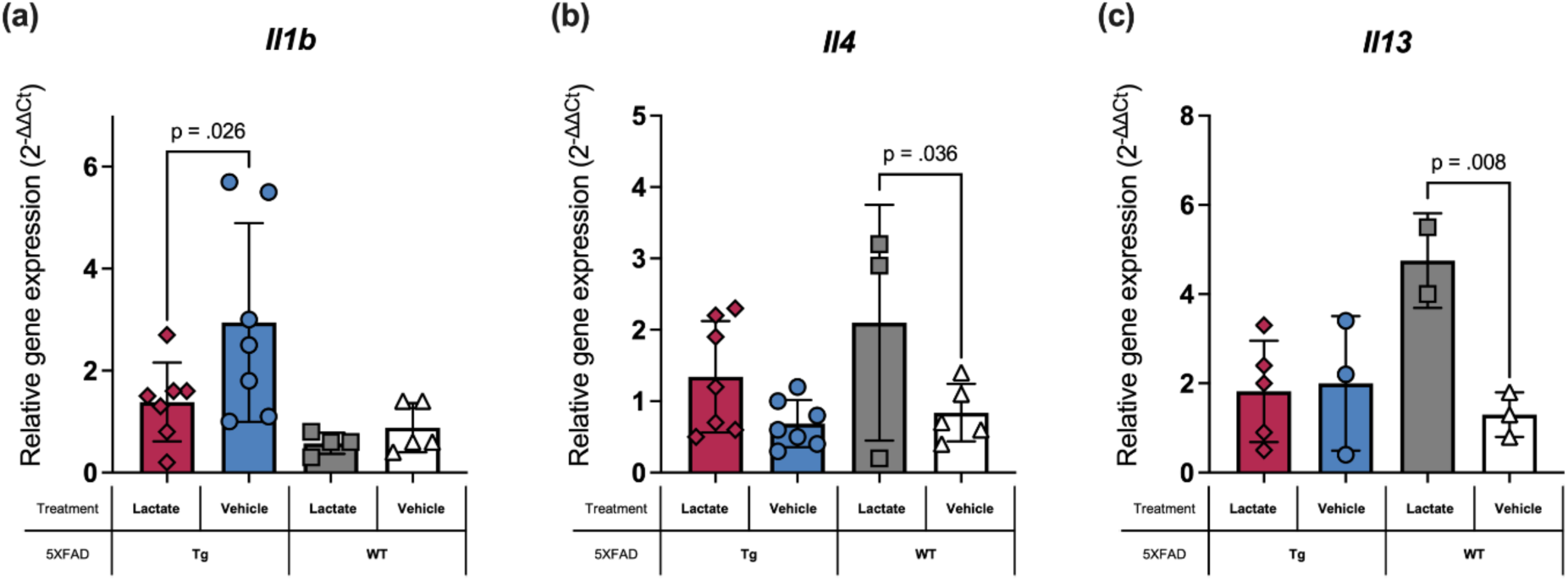
Early-lactate treatment reduces pro-inflammatory and enhances anti-inflammatory cytokines in 5XFAD transgenic (Tg) and wild-type (WT) mice. Early-lactate administration (1.1–2.3 months at the start, 4.0–5.0 months at the endpoint) reduced pro-inflammatory *Il1b* mRNA levels in Tg mice and increased anti-inflammatory *Il13* and *Il4* mRNA levels in WT mice, suggesting a protective effect. Relative gene expression was quantified by RT-qPCR using the 2^⁻ΔΔCt^ method. Data are presented as individual data points, with males and females combined. (a) *Il1b* mRNA was significantly downregulated in lactate-treated Tg mice (*n* = 7, *p* = 0.026, *d* = 1.30) compared to Tg controls (*n* = 7). (b) *Il4* mRNA was significantly upregulated in lactate-treated WT mice (*n* = 3) compared to WT vehicle controls (*n* = 5, *p* = 0.036, *d* = 1.66). A similar trend was observed in Tg mice, but the difference was not statistically significant. (c) *Il13* mRNA was significantly increased in lactate-treated WT mice (*n* = 2) compared to WT vehicle controls (*n* = 3, *p* = 0.008, *d* = 3.08). Statistical analyses were performed using one-way ANOVA followed by Fisher’s least significant difference test (LSD). Results are displayed as mean ± standard deviation (SD).

### Lactate treatment induces upregulation of lactate-handling protein mRNAs in WT mice

RT-qPCR analysis of hemibrains showed a modest, statistically non-significant increase of *Hcar1* in WT mice treated with lactate (Fig. 7a). However, lactate treatment significantly increased monocarboxylate transporter 2 (*Mct2*, *Slc16a7*) mRNA in WT mice (*p* = 0.021; Fig. 7b). Stratified analyses indicated a significant effect in WT females (*p* = 0.009), but not WT males (Supplementary Fig. 10a,b), and in late-treated WT mice (*p* = 0.032), but not early-treated WT mice (Supplementary Fig. 10c,d). Both *Hcar1* and *Mct2* were detected by RNA-seq; notably, *Hcar1* trended upward in lactate-treated mice relative to vehicle-treated (Supplementary Fig. 11).

**Figure 7.**
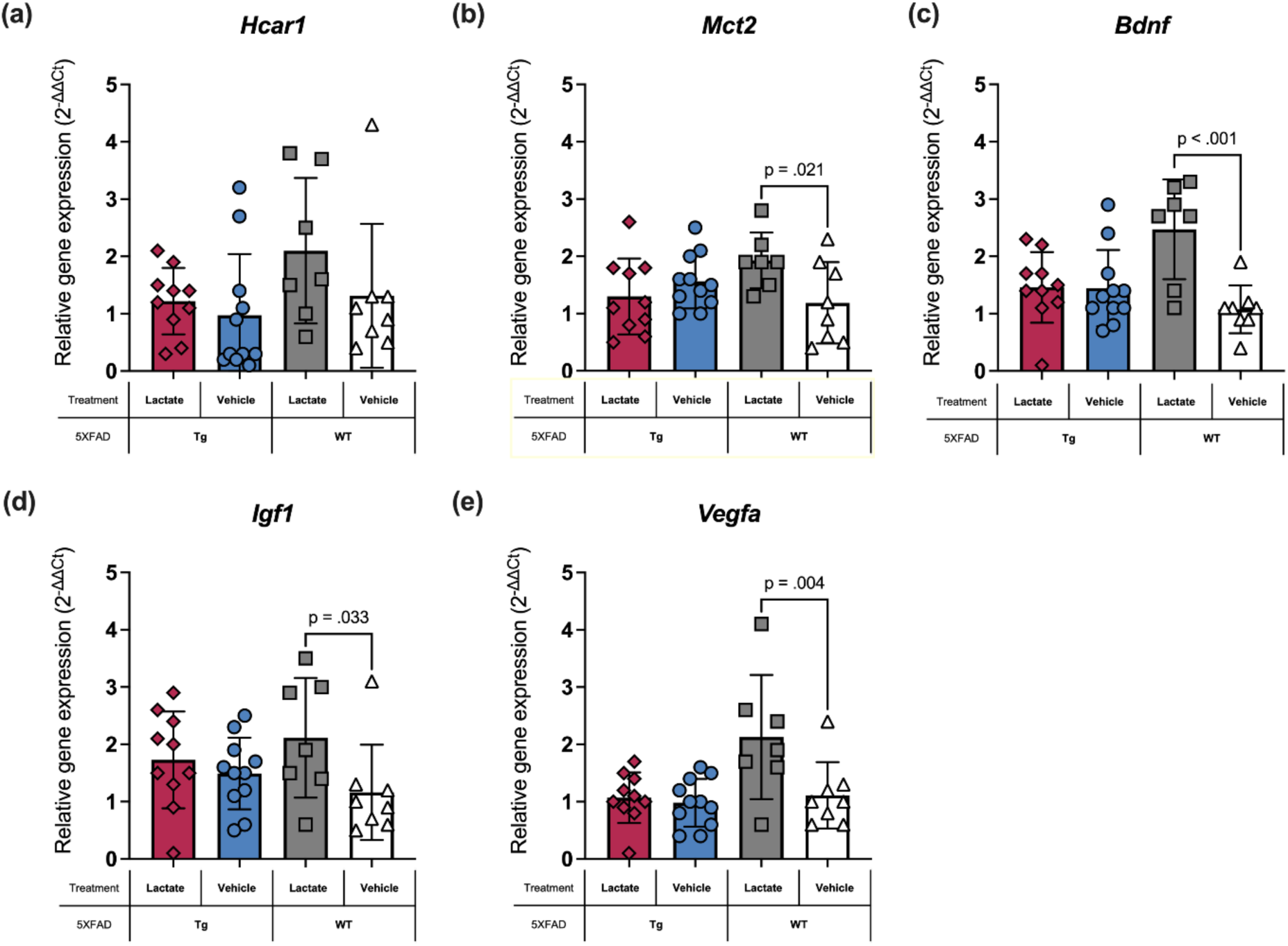
mRNA expression of receptor (*Hcar1*), transporter (*Mct2, Slc16a7*), and growth factors (*Bdnf, Igf1, Vegfa*) following lactate treatment. Relative mRNA expression levels of hydroxycarboxylic acid receptor 1 (*Hcar1*), monocarboxylate transporter 2 (*Mct2, Slc16a7*), Vascular endothelial growth factor (*Vegfa*), Brain-derived neurotrophic factor (*Bdnf*), and Insulin-like growth factor 1 (*Igf1*) were quantified by RT-qPCR using the 2^⁻ΔΔCt^ method in 5XFAD transgenic (Tg) and wild-type (WT) mice treated with lactate or vehicle. Tg lactate (*n* = 10), Tg vehicle (*n* = 11), WT lactate (*n* = 7), and WT vehicle (*n* = 8). Groups included both sexes and combined early- and late-treatment start cohorts. (a) *Hcar1* expression showed a modest increase following lactate treatment in WT mice; however, no significant differences were observed in any of the groups. **(b)** Lactate administration significantly increased *Mct2* expression in WT mice compared to vehicle-treated controls (*p* = 0.021, *d* = 1.26). No significant changes were observed in Tg mice treated with lactate versus vehicle. (c) *Bdnf* expression was significantly higher in WT mice receiving lactate compared to vehicle-treated WT controls (*p* < 0.001, *d* = 2.14). (d) *Igf1* expression was significantly upregulated in WT mice receiving lactate compared to WT controls (*p* = 0.033, *d* = 1.16). (e) *Vegfa* expression was significantly upregulated in lactate-treated WT mice compared to vehicle-treated WT controls (*p* = 0.004, *d* = 1.60). Data are presented as individual data points, with statistical analyses performed using one-way ANOVA, followed by Fisher’s least significant test (LSD). Results are displayed as mean ± standard deviation (SD).

### Lactate treatment enhances growth factor expression in WT mice

RT-qPCR of hemibrains revealed increased expression of the growth factors *Bdnf* (*p* < 0.001), *Igf1* (*p* = 0.033) and *Vegfa* (*p* = 0.004) in WT mice after lactate treatment (Fig. 7c–e). In Tg mice, no significant changes were observed. These transcripts were not detected by RNA-seq in our dataset. Together these data indicate that lactate enhances neurotrophic gene expression in WT brain, whereas the effect was not evident in Tg mice, consistent with disease-related blunting of transcriptional responsiveness.

## Discussion

Despite accumulating evidence that exercise-derived lactate contributes to brain plasticity and resilience, its direct and chronic effects in AD models have remained largely unexplored. Here, we demonstrate for the first time that sustained systemic lactate administration mitigates working memory decline in 5XFAD mice and induces widespread ameliorative molecular adaptations across synaptic, cytoskeletal, inflammatory, mitochondrial, and proteostatic pathways.

A recent study in 3xTg-AD mice, published after the completion of the present work, reported similar findings^18^, showing that IP lactate injections improved spatial working memory, increased synaptic density, and elevated hippocampal lactate levels. These findings align with previous work in WT mice, showing that astrocyte-derived lactate is required for long-term potentiation (LTP)^19,20^; that intrahippocampal lactate infusion improves spatial memory via neuronal MCT2^21^, which is enriched at the postsynaptic density of hippocampal neurons^22,23^; and that oral lactate administration combined with mild exercise enhances Y-maze performance^24^. Together, these findings position lactate as an enhancer of memory processes across health and AD.

Mechanistically, lactate acts as both fuel and signal. It crosses the blood-brain barrier supporting neuronal energy metabolism^25^; contributing to lipid synthesis^26^, replenishing tricarboxylic acid cycle intermediates^27^ and modulating redox via the NADH/NAD^+^ ratio^28^; promoting protein lactylation^29^ linked to adult hippocampal neurogenesis^30^; and providing the N-methyl-D-aspartate (NMDA) receptor co-agonist D-serine^31^. It also signals through HCAR1^7,8,32^, potentially regulating a disintegrin and metalloproteinase domain-containing protein 10 (ADAM10) activity and reducing Aβ by altering APP processing^33^. Lactate arising from aerobic glycolysis, i.e. the Warburg effect, may further drive synaptic growth and remodelling^34^. Together, these mechanisms link lactate metabolism and signalling to structural plasticity, neuroprotection, and amyloid regulation.

No available animal model fully reproduces human AD pathology. The 5XFAD mouse model^12^ expresses five familial AD mutations in APP and presenilin-1 (PSEN1), leading to early intraneuronal Aβ_42_ accumulation (from ∼1.5 months), extracellular plaque deposition (from 2 months), gliosis, neuroinflammation, synaptic and neuronal loss, and memory impairment (from 4-6 months) (Fig. 1b). Phosphorylated tau aggregates have also been reported^35^. Although based on artificial overexpression, 5XFAD recapitulates key pathological features of human AD and is therefore appropriate for initial testing of lactate interventions.

Lactate treatment produced peak blood levels within the physiological range of high-intensity exercise^7^, and below those seen during extreme voluntary exertion in humans (∼32 mmol L⁻¹)^36^. To minimise irritation and stress, we alternated IP and SC injections. IP delivery yields rapid, transient peaks via first-pass hepatic metabolism, whereas SC absorption is slower, but circumvents an initial liver pass; nonetheless, prior work^10^ indicates broadly comparable systemic kinetics. Accordingly, daily lactate injections generated transient physiological surges (peak within a quarter of an hour, normalization within one hour) rather than sustained hyperlactataemia. Treatment was well tolerated, with no anxiety-like behaviour in the open-field test, in contrast to reports of panic in susceptible human subjects^13^ and anxiety-like responses in rodents^14^, suggesting that anxiogenic effects are unlikely to limit clinical application.

Lactate strengthened synaptic robustness. NEFL, a cytoskeletal protein elevated in AD biofluids^37^, was increased in Tg mice, but reduced by lactate in Tg females, consistent with neuroprotection (Fig. 4a). In contrast, WT females showed increased NEFL with lactate, possibly suggesting cytoskeletal remodelling in the healthy brain. Similarly, in WT males, lactate upregulated *Nefh,* which encodes the heavy neurofilament subunit essential for axonal structure, transport, and plasticity^38^ (Fig. 5e). These findings indicate that lactate exerts context-dependent, bidirectional effects, enhancing structural resilience under pathological conditions while supporting remodelling in the healthy brain.

OPCML, an IgLON family adhesion molecule that shapes synaptic organisation and is typically reduced in AD, was restored in Tg males with lactate (Fig. 4b). This supports a role for lactate in reinforcing adhesion and structural stability at the synapse. PPFIA2, a presynaptic active-zone scaffold that organises vesicle release, was upregulated in Tg females (Fig. 4c). Together with STXBP3, a syntaxin-binding protein required for SNARE-mediated vesicle fusion, and SYT1, the Ca²⁺ sensor for fast vesicle release, both of which were restored in Tg males (Fig. 4d,e), these changes suggest that lactate strengthens presynaptic architecture and neurotransmitter release capacity. VGLUT2, the vesicular glutamate transporter required for glutamate loading into synaptic vesicles, was also increased by lactate in WT females, WT males, and Tg males, whereas vehicle-treated Tg mice showed reduced expression compared to WT (Fig. 4f). Beyond its canonical role, VGLUT2 functions as a “differentiation-associated Na⁺-dependent inorganic phosphate transporter” (DNPI)^39^, highlighting a dual role in development^15,16^ and metabolism that may be compromised during neurodegeneration. Consistent with this, VGLUT2 expression declines with ageing^40^ and is reduced in post-mortem AD brain regions^41^. Misfolded tau also accumulates at VGLUT2-positive terminals in the anterodorsal thalamic nucleus^42^, in line with the selective vulnerability of these synapses. These findings underscore the central role of glutamate as the brain’s principal excitatory neurotransmitter. Accordingly, ionotropic receptor subunits are reduced in AD^43^, whereas lactate upregulated ionotropic glutamate receptor transcripts from low levels in Tg mice (Fig. 5f–h). Thus, the lactate-induced increase in VGLUT2 may contribute to preserving synaptic integrity. Finally, VSNL1, a neuronal Ca²⁺ sensor of the NCS family and established AD biomarker, was upregulated in Tg females (Fig. 4g). In clinical studies, elevated VSNL1 in cerebrospinal fluid correlates with tau pathology, brain atrophy, and cognitive decline^44^. However, in our experimental context — where lactate improved working memory and enhanced synaptic protein expression — VSNL1 upregulation may instead reflect adaptive Ca²⁺-dependent signalling and neuronal resilience. This suggests that lactate not only restores presynaptic machinery but may also reframe molecular markers of degeneration as indicators of strengthened synaptic function.

In the perisynaptic space, HAPLN4, a hyaluronan–proteoglycan linker that stabilises perineuronal nets and nodes of Ranvier, was increased in WT males (Fig. 3d; Supplementary Fig. 7f). At the postsynaptic density, NTRK2, the TrkB receptor for BDNF, which is critical for synaptic plasticity and neuronal survival, was upregulated by lactate in WT females (Fig. 3c; Supplementary Fig. 7g). Together, these adaptations suggest that lactate enhances extracellular matrix stability and postsynaptic plasticity signalling in the healthy brain.

Core AD-associated proteins, APP, APOE, CLU and GFAP, were elevated in Tg mice (Fig. 3c; Supplementary Fig. 7a–d), consistent with gliosis and neuroinflammation in the AD phenotype, processes recognised as central drivers of AD pathology^45^. Lactate did not significantly alter these markers, although levels in lactate-treated Tg mice trended towards WT. Phospholipase D family member 3 (PLD3), an endolysosomal AD risk factor, showed a similar trending pattern, but no statistically significant change with lactate (Supplementary Fig. 7e). At the transcript level, lactate reduced the pro-inflammatory cytokine interleukin-1b (*Il1b*) in Tg mice (Fig. 6a) and showed increases of anti-inflammatory interleukin-4 (*Il4*) and interleukin-13 (*Il13*) in WT mice (Fig. 6b,c). These findings indicate a modest protective inflammation- and gliosis-attenuating effect of lactate, complementing its synaptic actions without directly altering core amyloid/glial proteins in the AD mice.

Lactate modulated proteins involved in mitochondrial energetics and cellular metabolism in a genotype- and sex-dependent manner. ATP5G2, increased in Tg females, indicating enhanced oxidative phosphorylation; GrpE-like 1, mitochondrial (GRPEL1), and solute carrier family 25 member 23 (SLC25A23), were upregulated in WT males, consistent with strengthened mitochondrial protein import and Ca²⁺-regulated ATP-Mg/Pi transport (Fig. 3c; Supplementary Fig. 7h–j). Apurinic/apyrimidinic endonuclease 1 (APEX1), increased by lactate in WT females; farnesyl diphosphate synthase (FDPS), was elevated in Tg groups irrespective of lactate treatment; GDP-mannose pyrophosphorylase B (GMPPB), decreased in WT males; guanylate kinase 1 (GUK1), decreased in Tg males with lactate; and 5-oxoprolinase (OPLAH), was downregulated in WT males, consistent with reinforcement of glutathione-cycle redox control (Fig. 3c; Supplementary Fig. 7k–o). Collectively, these adaptations suggest that lactate augments mitochondrial energetics and import while tuning nucleotide, sterol, and glutathione-linked redox metabolism — changes that plausibly support synaptic resilience and the behavioural benefits observed in Tg mice.

Sex-stratified analyses revealed marked variability in both baseline differences and lactate responses, consistent with the higher prevalence of AD in women and the recognised influence of sex hormones and chromosomes on disease mechanisms^46^. In females, lactate trended to reduce AD markers (Fig. 3c; Supplementary Fig. 7a–e), increased ionotropic glutamate receptor transcripts in Tg mice (*Grik3, Grin2c, Grid2ip*; Fig. 5f–h), and elevated ATP5G2 in Tg animals (Fig. 3c; Supplementary Fig. 7h), consistent with known sex-dependent regulation^47^. *Grik3* encodes a kainate receptor subunit that can heteromerise with GRIK4 or GRIK5 to form excitatory receptors, supporting synaptic transmission across the central nervous system. *Grin2c* encodes a subunit of the NMDA-receptor, which plays a pivotal role in synaptic plasticity and memory, and is critical for LTP induction^48^. Notably, NMDA receptors require glycine or D-serine as a co-transmitter to be sensitive to glutamate^49^, which is particularly interesting given recent evidence that lactate increases astrocytic formation and release of D-serine via HCAR1^31^. *Grid2ip*, encoding glutamate receptor delta-2 interacting protein, serves as a postsynaptic scaffold linking glutamate receptor subunit δ2 (GRID2) to intracellular signalling complexes and is essential for synapse organisation and plasticity in Purkinje cells. Together, these findings indicate that lactate enhanced postsynaptic responsiveness in Tg females.

In males, lactate downregulated pro-apoptotic regulators: arginine–glutamic acid dipeptide repeats (*Rere*) and histone deacetylase 4 (*Hdac4)* (Supplementary Fig. 6c,d), implying reduced apoptotic drive and de-repression of myocyte enhancer factor 2C/D (MEF2C/D) mediated transcription. Lactate also upregulated mitochondrial proteins GRPEL1 and SLC25A23 (Fig. 3c; Supplementary Fig. 7i,j), enhancing mitochondrial protein import and energetics, while downregulating methylenetetrahydrofolate reductase (*Mthfr*), a key enzyme in folate metabolism whose polymorphisms are linked to elevated homocysteine levels and increased AD risk^50^ (Fig. 5e).

In Tg females, lactate upregulated autophagy-related 4A cysteine peptidase (*Atg4a*), essential for autophagosome formation^51^. This is in line with the view that impaired autophagy contributes to AD pathogenesis^52^ and the reported ATG4A dysregulation in neurons differentiated from AD patient derived induced pluripotent stem cells^53^. Lactate also increased ubiquitin protein ligase E3 component N-recognin 2 (*Ubr2*), implicated in the ubiquitination of pathological AD markers^54^, consistent with lactate-reinforced autophagy and proteostasis. (Fig. 5b).

Several lactate-induced changes were observed only in WT mice, including upregulation of *Mct2, Bdnf, Igf1, Vegfa,* and regulations of mitochondrial proteins GRPEL1 and SLC25A23, suggesting that AD pathology may blunt some of lactate’s actions and highlighting a preventive potential. Taken together, lactate engages divergent sex-specific pathways strengthening glutamatergic signalling and autophagy in females, while modulating transcriptional plasticity, mitochondrial energetics, and folate metabolism in males. These findings underscore sex as a critical biological variable and highlights the need for precision-medicine approaches in AD.

This exploratory study has several limitations that should be considered. Use of hemibrains facilitated detection of broad molecular changes but may have masked region-selective effects. Group sizes were small in some assays, particularly for RT-qPCR data on *Il4* and *Il13* in WT mice (*n* = 2-3), limiting statistical power. Classical histopathological endpoints (e.g. Aβ plaque burden, microglial morphology, synaptic density) were not assessed, restricting conclusions on disease modification. The timing of treatment initiation may also be a confounder: in early-treatment cohorts, pathology was minimal, and lactate may have acted preventively, but the observation time may have been too short to find the full effect on chemistry and behaviour. In the late-treatment cohort, Aβ deposition and gliosis were already emerging, potentially reducing efficacy. Systemic tolerability markers were not measured, though stable body weight and absence of anxiety-like behaviour indicated satisfactory tolerability. Oxidative stress markers were also lacking, leaving open whether lactate-induced mitochondrial activation has net protective or adverse redox effects. Finally, brain lactate was not measured directly, although *Mct2* upregulation and prior evidence of blood-brain barrier transport support central exposure. Behavioural assessment was restricted to the Y-maze and additional paradigms will be needed to determine whether effects generalise across cognitive domains.

In conclusion, this study demonstrates that repeated lactate administration mitigates working memory decline in 5XFAD mice and beneficially modulates multiple molecular pathways associated with AD. Lactate enhanced synaptic proteins (OPCML, PPFIA2, STXBP3, SYT1, VGLUT2, and VSNL1), attenuated neuronal injury markers (NEFL), reduced pro-inflammatory signalling (*Il1b*), and supported mitochondrial and proteostatic factors. A particularly striking finding was the restoration of glutamatergic markers typically reduced in AD, pointing to a mechanism whereby lactate may strengthen synaptic efficacy and neuronal resilience. Importantly, both lactate’s effects and the baseline Tg-WT differences varied by sex, underscoring the need for precision medicine approaches. These findings provide convergent behavioural, proteomic, and transcriptomic evidence of lactate’s benefits, encouraging further mechanistic and translational studies, in larger, age-stratified cohorts with extended behavioural paradigms to better define lactate’s therapeutic and preventive potential in AD.

## Materials and methods

### Mice

Animal experiments were conducted in accordance with national and regional ethical guidelines at the Institute of Basic Medical Sciences, Faculty of Medicine, University of Oslo and complied with the ARRIVE guidelines (https://arriveguidelines.org). All animal procedures were approved by the Norwegian Food Safety Authority and the Norwegian Animal Research Authority (FOTS 13167, 15525), adhering to the standards set by the European Union.

Mice were group-housed in Seal Safe Plus GM900 and GM500 cages (Scanbur AS, Norway), with woodchip bedding, nesting paper, and in-cage shelters. Mice were regularly monitored for signs of aggression and those exhibiting such signs were rehoused in smaller groups or individually, as needed, to ensure their welfare. All mice had *ad libitum* access to water and a standard pellet diet (Teklad Global 18% Protein Rodent Diet (2018S), Envigo, UK). The housing environment was maintained at 22 ± 1 °C with 55 ± 5% humidity, and mice were kept on a 12-h light/dark cycle (7 am to 7 pm). Treatments and experiments were performed during the light phase (9 am to 5 pm), corresponding to the mice’s inactive/resting period. Light intensity ranged from 45 lux during the light cycle to 285 lux during peak illumination.

5XFAD B6/SJL mice were acquired from The Jackson Laboratory (34840-JAX, USA) and bred at the Institute of Basic Medical Sciences, University of Oslo. The 5XFAD mouse model overexpresses human *APP* and *PSEN1* transgenes carrying five AD mutations (including Swedish, Florida, and London mutations in *App* and M146L and L286V in *Psen1*) under the control of the thymus cell antigen 1 (*Thy1*) promoter, enabling targeted expression in nervous system cells.

The AD mice used in this study were hemizygous for both *App* and *Psen1* transgenes, generated from the 5XFAD model, along with WT littermate controls. Of note, the *App* transgene includes the 5’ untranslated region, which contains a putative interleukin-1b (*Il1b*) translational enhancer element.

Genotyping was conducted on 6–8-week-old mice using ear-punch biopsy samples. DNA was isolated using the Extract-N-Amp Tissue PCR kit (XNAT2; Sigma Aldrich, USA), followed by PCR and agarose gel electrophoresis to identify the presence of *App* and *Psen1* transgenes (Supplementary Table 1). The experimental design was carefully structured to ensure balanced representation of male and female mice.

### Treatments

Treatments were initiated at two distinct age intervals: an early-treatment start (1.1–2.3 months old), and a late-treatment start (2.6–3.2 months old) (Fig. 1b). Within each age group, mice were randomly assigned to either the vehicle group (1× PBS; 10010023, ThermoFisher, UK) or the lactate group (Sodium L-lactate; 71718, Sigma-Aldrich, MO, USA) (Fig. 1a). Mice in both groups received treatments four times per week for 11 weeks using 12.7 mm/29-gauge syringes (BD Micro-Fine+ 1 mL Insulin). Mice in the lactate group were administered 2 g kg⁻¹ body weight of sodium L-lactate (18 mmol kg⁻¹) dissolved in 1× PBS (pH 7.4) at 100 mg mL⁻¹. This dose was chosen to replicate lactate levels observed after high-intensity training^7^ and was adjusted for long-term administration (Supplementary Fig. 1). To maintain treatment efficacy while minimising localised tissue reactions, injection routes alternated between SC and IP^10^.

### Behavioural tests

OFT was used to assess anxiety-like behaviour following lactate injections^13^, while the Y-maze was employed to evaluate working memory^55^. Behavioural data from both tests were analysed using BiKiPy, a Python package specifically designed for behavioural analysis. Mouse coordinates were determined using DeepLabCut, and the resulting data were smoothed using a Seasonal Autoregressive Integrated Moving Average with Exogenous factors (SARIMAX) model to improve signal clarity and reduce noise.

### OFT

To assess anxiety-like behaviour, mice were observed for 5 min as they explored a novel, roofless square arena measuring 40 x 40 x 34 cm, illuminated at 15 lux. The arena was divided into two distinct zones: a central zone (20 x 20 cm) and a peripheral zone (a 10 cm band along the arena’s edges). To prevent confounding odours, the arena was cleaned with 70% ethanol between trials. Behavioural data were recorded using ANY-maze tracking software (v6.21, Stoelting Co., USA).

For analyses, confinement in the centre across video frames was quantified by defining the centre as a rectangular perimeter and the periphery as the coordinates outside this perimeter. Confinement within the rectangular perimeter was subjected to tolerance modelling, yielding centre confinement (CC). By flipping the binary values and applying tolerance modelling, periphery confinement (PC) was obtained. CC and PC are binary sequences in which a value of 1 indicates confinement to the respective zone. To calculate the time spent in each zone, the values in the corresponding sequences are summed and the result divided by the frame rate (frames per second; fps). This provides the total duration, in seconds, that the animal spends in each area.

### Y-maze test

The Y-maze consisted of three identical arms (35 x 8 x 45 cm) arranged at 120° angles. The test was conducted in low-light conditions, with an 8-lux light source positioned above the maze. Prior to the test, mice were acclimatised to the testing room for 30 min. During the 6-min trial, each mouse was placed in one arm of the maze and allowed to explore freely. An arm entry or exit was considered complete when the centre of the animal’s body crossed the arm entrance. To control for odour cues, the maze was thoroughly cleaned with 70% ethanol between trials.

For analyses, a reduced version of the original dataset that contains the location of the animal in every recorded frame was used. This reduction eliminates redundant data by compressing consecutive identical locations into single entries. A tolerance model was applied to consecutive arm entries. Area alternations were calculated as the count of each area label in the dataset, with the results stored in a map (key: area, value: alternations). When a mouse visits three unique arms consecutively, it’s called a triple alternation. The sum of triplet alternations is the length of the area alternation sequence minus two. The subtraction is used to ignore length of the two initial elements from the reduced sequence.

Spontaneous alternations (SA) are computed as follows:

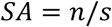

Where *n* is the total number of unique triplet alternations and *s* is the sum of total alternations.

Distance travelled (displacement): To get the frame-wise displacement the coordinates have their origin-distance computed by the Euclidean norm. The finite difference of the origin-distance gives frame-wise displacement:

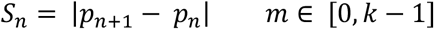

The total displacement is computed by:

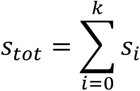

Where *p* is position; *s* is displacement; *k* is the number of frames in the video recording.

### Brain tissue preparation

Following the completion of behavioural testing, mice were euthanised via cervical dislocation. Brains were quickly dissected and hemisected. The whole-brain hemispheres were immediately frozen by placement on solid CO_2_ and subsequently stored at -80°C for homogenization and extraction, as described below.

### Protein sample preparation and LC-MS/MS analysis

Protein pellets were precipitated using the AllPrep DNA/RNA/Protein Mini Kit (80004; Qiagen), and protein concentrations were determined via the BCA assay (Pierce). For each replicate, 20 μg of protein was precipitated on amine beads, following a previously described method^56^.

Immobilised proteins on beads were dissolved in 50 mM ammonium bicarbonate, reduced, alkylated, and digested with trypsin (1:50 enzyme ratio; Promega) at 37 °C overnight.

Peptides were loaded onto Evotips and analysed by LC-MS/MS using the Evosep One system (Evosep), coupled with a Q Exactive HF mass spectrometer (ThermoFisher). The analysis employed an 88-min gradient (Evosep extended method).

Raw data from the LC-MS/MS analysis were processed using MaxQuant software (v 1.6.17.0) for protein identification and label-free quantification. Carbamidomethyl (C) was set as a fixed modification, while N-acetylation and methionine oxidation were variable modifications. The peptide search was configured with a first search tolerance of 20 ppm and a main search tolerance of 4.5 ppm. Trypsin (without proline restriction) was selected as the enzyme, allowing up to two missed cleavages. The minimum requirement for unique peptides was set to one, and the false discovery rate (FDR) was set at 1% (FDR ≤ 0.01) for both peptide and protein identification. The UniProt mouse database was employed, and reversed sequences were generated for FDR rate assignment. Known contaminants and reversed entries, as provided by MaxQuant and identified in the samples, were excluded from further analysis.

The “proteinGroups.txt” file containing quantified protein data was used for downstream analysis, mainly including differential expression analysis using the DEP R package (v 1.12.0). Briefly, the data was cleaned by removing contaminants and reverse proteins, and proteins missing label-free quantification (LFQ intensity) values in all replicates in at least one condition. Data normalisation used variance stabilizing transformation (vsn) and a limma-based model. Missing values were imputed using MinProb, suitable for “missing not at random” (MNAR) data and random selection of minimal values from a left-shifted distribution. DEPs were identified using the protein-wise linear model (limma package within DEP package) combined with empirical Bayes statistics, with an adjusted *p*-value < 0.05 and a minimum absolute log_2_ fold change > 0.5. Gene ontology (GO) and Kyoto Encyclopedia of Genes and Genomes (KEGG) pathway enrichment analyses were conducted using the clusterProfiler R package (v 3.18.1).

### RNA-seq analysis

mRNA was enriched from total RNA samples using Oligo(dT) magnetic beads, followed by conversion to cDNA via reverse transcription with an N6 primer. Adaptors were ligated to the cDNA ends, and ligation products were amplified through PCR using two specific primers. The PCR products were denatured, and single-stranded DNA (ssDNA) was cyclised using splint oligos and DNA ligase. The library was prepared and sequenced on an SE50 instrument at BGI (Hong Kong, China). Raw reads underwent quality control (QC) and were filtered into clean reads, which were aligned to the mm10/GRCm38 mouse reference genome using HISAT^57^ (index file: https://cloud.biohpc.swmed.edu/index.php/s/grcm38_tran/download<u>; Gtf file:</u> http://hgdownload.soe.ucsc.edu/goldenPath/mm10/bigZips/genes/mm10.refGene.gtf.gz).

The reads alignment and transcripts assembly were performed on a Saga clustering computer provided by UNINETT Sigma2. Aligned SAM files were sorted and converted to BAM files using SAMtools. Transcripts were assembled with stringTie (v1.3.5) and merged using the stringTie –merge function. Read coverage tables with normalised Fragments Per Kilobase of transcript per Million mapped reads (FPKM) counts were generated based on merged transcripts using the stringTie –eB function. DESeq2 (R package, v1.30.1) was employed to compare transcripts, normalise counts, and identify DEGs. Raw counts were normalised by size factor, which is the median of ratios calculated by dividing the raw count with the geometric mean of each gene among all samples. The differential expression was tested using negative binomial generalised linear models. A modified t-test (Wald test) was applied for *p*-value analysis, and adjusted *p*-values were obtained using the Benjamini-Hochberg (BH) procedure. DEGs were determined as absolute log_2_ fold change ≥ 1 and adjusted *p*-value < 0.05. DEGs were further processed and visualised with R packages, including ggplot2, tidyverse, principal component analysis (PCA), ggvenn (v0.1.9, for Venn diagram), and pheatmap (v1.0.12). DEGs underwent Gene Ontology (GO) enrichment analysis and functional annotation with the R package clusterProfiler (v3.18.1)^58^. The overrepresentation module (enrichGO function) was used with an adjusted *p*-value cutoff of 0.05 and pAdjustMethod set to “BH”. Biological process (BP), molecular function (MF), and cellular component (CC) ontologies were analysed separately.

### RNA extraction and RT-qPCR

Snap-frozen mouse hemibrains were individually homogenised using a FastPrep-24 machine at a speed of 6.5 ms⁻¹ for 45 seconds in 1 mL of buffer RLT. Total RNA was extracted using the RNeasy Lipid Tissue Mini Kit (QIAGEN, Sweden) according to the manufacturer’s instructions and eluted in 30 μL of RNase-free water. RNA quality and concentration were assessed with a Nanodrop 2000c spectrophotometer (Thermo Scientific, USA), ensuring the samples were kept on ice during handling.

RNA concentrations were normalised to 38.2 ng/μL, and cDNA was synthesised using the Reverse Transcriptase Core Kit (Eurogentec, RT-RTCK-05; Lot. EUK211, Belgium) to obtain a final concentration of 1.09 ng/μL. Real-time quantitative PCR (RT-qPCR) was carried out using the Takyon low ROX Probe 2X MasterMix (dTTP blue - 750 rxn, UF-LPMT-B0701) and Taqman Gene Expression Assays (Thermo Fisher Scientific) on an Agilent Technologies AriaMx Real-Time PCR system. Genes of interest, including *Bdnf* (Mm01334042_m1), *Hcar1* (Mm00558586_s1), *Igf1* (Mm00439560_m1), *Il1b* (Mm00434228_m1), *Il4* (Mm00445259_m1), *Il13* (Mm00434204_m1), *Slc16a7 (Mct2)* (Mm00441442_m1), *Vegfa* (Mm00437306_m1), were assessed. The reference gene *Rpl13a* (Mm01612986_gH) was used for normalisation, and relative expression levels were calculated using the ΔΔCt method.

### Statistical analysis

All individual measurements constitute biological replicates. Sex was included as a biological variable in all analyses, and females and males were evaluated separately to assess potential sex-dependent effects. Groups with n < 9 were analysed using parametric tests. Groups with n ≥ 9 were first evaluated for normal distribution using D’Agostino and Pearson’s omnibus normality test. The specific statistical approaches applied to each experiment are detailed in the figure legends. Data are presented as box-and-whisker plots (median and interquartile range) or as bar plots (mean ± standard deviation or standard error of the mean). Statistical analyses were performed using GraphPad Prism (v10.2.0; GraphPad Software). Given the factorial design of the behavioural experiments, data were analysed by two-way ANOVA followed by Tukey’s post hoc test to assess group differences and potential interactions. For molecular assays (RT-qPCR and a subset of proteomics), analyses were performed separately within each sex using one-way ANOVA followed by Tukey’s test or Fisher’s least significant test, due to smaller group sizes and partial factorial design. For RNA-seq and Proteomics, differential expression was assessed using Wald statistics with Benjamini-Hochberg correction to control the false discovery rate. Effect sizes are reported as Cohen’s *d* for post hoc comparisons and η² (partial η²) for ANOVA. Statistical significance was set at *p* < 0.05 and is indicated as follows: \**p* < 0.05, \*\**p* < 0.01 and \*\*\**p* < 0.001.

## Supporting information

Supplementary_Figures_and Table_Belhaj_Amellem

## Acknowledgements

The proteomic analyses, employing mass spectrometry, were conducted at the Proteomics Core Facility, Department of Immunology, University of Oslo/Oslo University Hospital. We would like to acknowledge the support received from the Core Facilities program of the South-Eastern Norway Regional Health Authority, which has greatly contributed to the success of this research endeavour. We are also grateful for the affiliation of this core facility with the National Network of Advanced Proteomics Infrastructure (NAPI), funded by the Research Council of Norway’s INFRASTRUKTUR-program (project number: 295910).

## Author contributions

IB, IÅ, JSM, and LB contributed to the conception and design of the study. IB, IÅ, HMW, and EMV conducted the experiments. IB and IÅ were responsible for data collection and analysis. CHT developed the behavioural data analysis software, while MY performed the RNA-seq and proteomic data analyses. MB provided funding for the study, and FAC supplied the mouse line. IB, IÅ, JSM, and LB co-wrote and revised the manuscript. All authors reviewed and approved the final version of the manuscript.

## Data availability statement

The RNA-seq data generated in this study have been deposited in the Gene Expression Omnibus (GEO) under the accession code GSE276471. Additional data supporting the findings of this study are available from the corresponding author upon reasonable request.

## Additional information

Supplementary information.

