## Supplementary_Figures_and Table_Belhaj_Amellem for "Lactate treatment improves brain biochemistry and cognitive function in transgenic Alzheimer’s and wild-type mice"

[1] Brain and Muscle Energy Group, Institute of Oral Biology, University of Oslo, Oslo, Norway.
[2] Centre for Embryology and Healthy Development (CRESCO), University of Oslo, Oslo, 0373, Norway.
[3] Department of Clinical and Molecular Medicine, Norwegian University of Science and Technology, Trondheim, Norway.
[4] Department of Neurosciences, Charité University Hospital, The German Center for Neurodegenerative Diseases (DZNE), Berlin, Germany.
[5] Department of Medical Biochemistry and Department of Microbiology, Oslo University Hospital and University of Oslo, Oslo, Norway.
[6] Division of Anatomy, Department of Molecular Medicine, Institute of Basic Medical Sciences, University of Oslo, Oslo, Norway.
[7] Department of Pathology, Oslo University Hospital, Norway
[8] KAUST Center of Excellence for Smart Health, King Abdullah University of Science and Technology 23955, Saudi Arabia.
† These authors contributed equally to this work and share first authorship

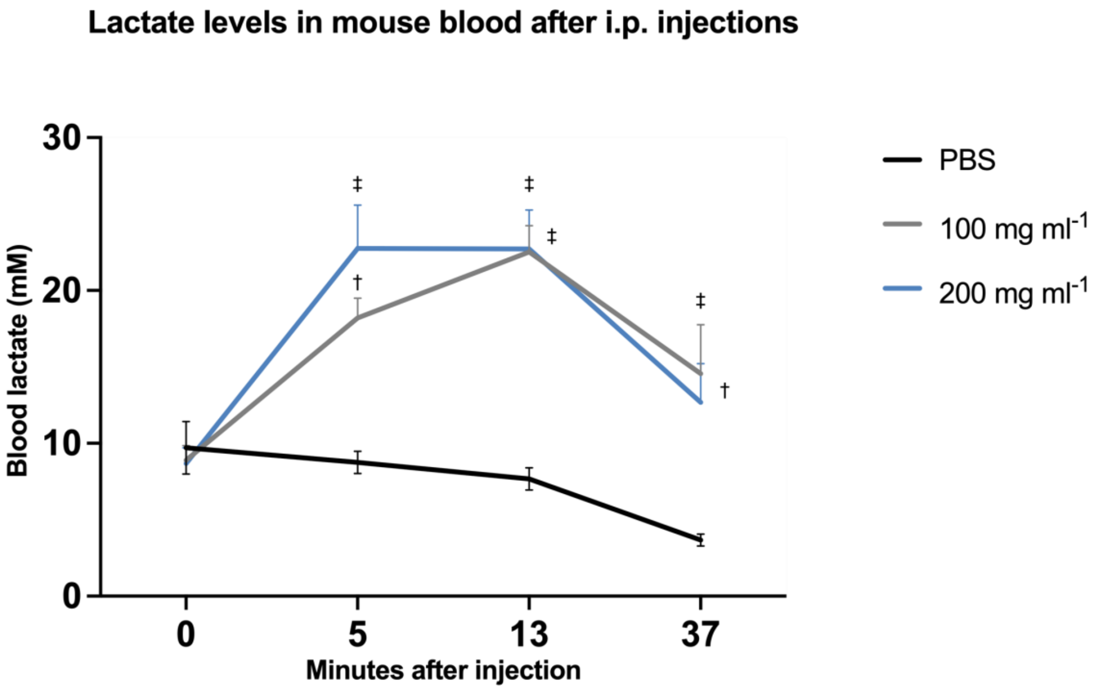


**Supplementary Figure 1. Blood lactate levels following intraperitoneal (IP) injections.** Blood lactate levels were measured in 5XFAD mice following IP administration of sodium L-lactate dissolved in 1× phosphate buffered saline (PBS) (pH 7.4) or 1× PBS alone in a separate cohort of mice prior to the experimental study. Sodium L-lactate was injected at a dose of 2 g $kg^{-1}$ in two different concentrations: 100 mg $ml^{-1}$ ($n$ = 10) and 200 mg $ml^{-1}$ ($n$ = 9). Mean blood lactate levels peaked at 5 minutes post-injection, reaching 22.76 mM (‡) in the 200 mg $ml^{-1}$ group and 18.20 mM (†) in the 100 mg $ml^{-1}$ group. By 13 minutes post-injection, lactate levels were comparable between the two groups: 22.71 mM (‡) for 200 mg $ml^{-1}$ and 22.52 mM (‡) for 100 mg $ml^{-1}$. By 37 minutes, lactate levels declined to 12.69 mM (200 mg $ml^{-1}$) and 14.55 mM (100 mg $ml^{-1}$). No significant changes were observed in PBS-injected mice ($n$ = 9) compared to pre-injection levels. Time points with significantly higher blood lactate levels in lactate-treated mice versus PBS controls are indicated as follows: (†) $p < 0.01$ and (‡) $p < 0.001$ (Tukey's multiple comparisons test). Blood lactate levels (mM) are presented as mean ± standard error of the mean (SEM).

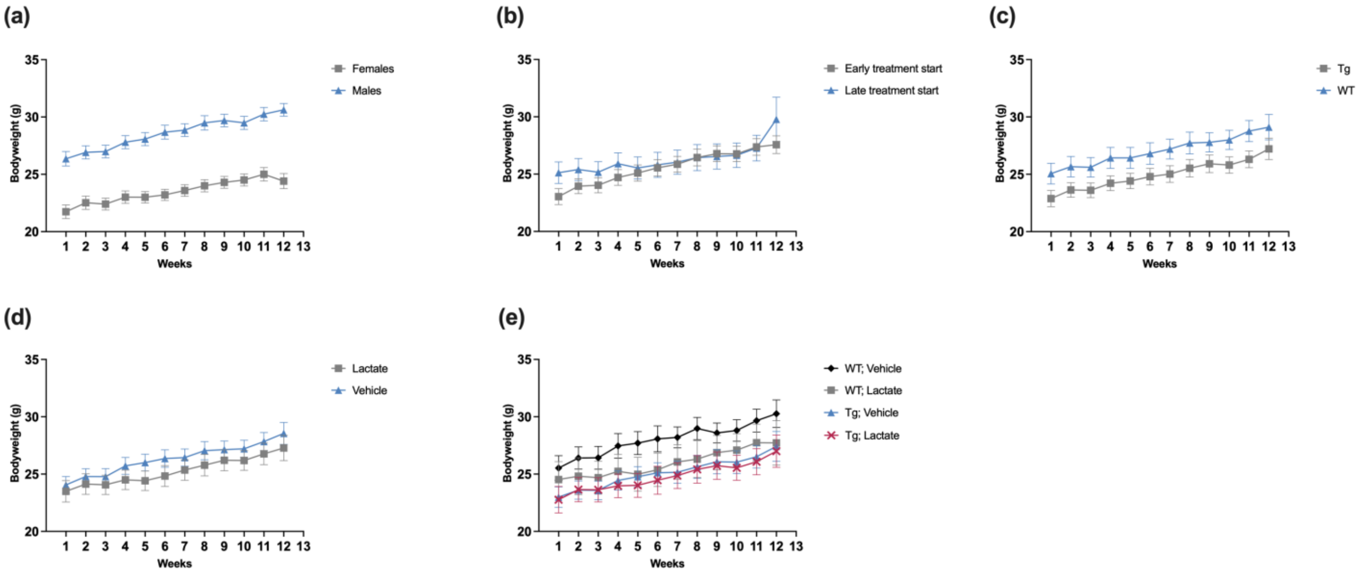


**Supplementary Figure 2. Body weight changes in 5XFAD transgenic (Tg) and wild-type (WT) mice during the treatment period.**
The graphs illustrate variations in mean body weight (g) over the treatment period in all 36 mice, with data presented across different group comparisons. **(a)** Comparisons between males (*n* = 16) and females (*n* = 20). **(b)** Comparisons between early-treatment start (1.1–2.3 months at the start, 4.0–5.0 months at the endpoint, *n* = 23) and late-treatment start (2.6–3.2 months at the start, 5.4–6.0 months at the endpoint, *n* = 13) groups. **(c)** Body weight changes in (WT, *n* = 15) and (Tg, *n* = 21) mice. **(d)** Comparisons between vehicle-treated (*n* = 19) and lactate-treated (*n* = 17) groups. **(e)** Body weight changes across the four experimental groups: WT Vehicle (*n* = 8), WT Lactate (*n* = 7), Tg Vehicle (*n* = 11), and Tg Lactate (*n* = 10). Bodyweight values are presented as g mean ± standard error of the mean (SEM).

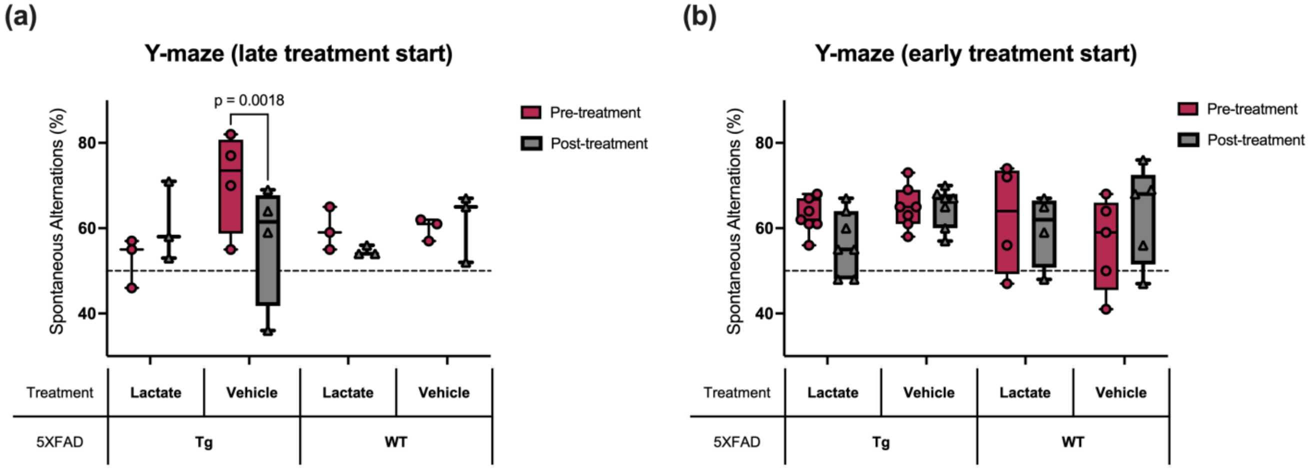


**Supplementary Figure 3. Raw spontaneous alternation (SA) percentages in the Y-maze in 5XFAD transgenic (Tg) and wild-type (WT) mice.**
Raw SA percentages before (Pre-treatment, red) and after (Post-treatment, grey) the treatment period are shown for the two treatment schedules. **(a)** Late-treatment start (2.6–3.2 months at the start; 5.4–6.0 months at the endpoint): Two-way repeated measures ANOVA revealed a significant Group × Time interaction. Tukey's post hoc tests confirmed a decline in Tg vehicle-treated mice ($p$ = 0.0018, $d$ = 3.10), whereas Tg lactate-treated mice were protected from decline. WT groups showed no significant pre–post changes. **(b)** Early-treatment start (1.1–2.3 months at the start; 4.0–5.0 months at the endpoint): No significant pre–post changes were detected across groups. Data are presented as box-and-whisker plots (median, interquartile range, all individual data points), with dashed lines indicating chance level (50%).

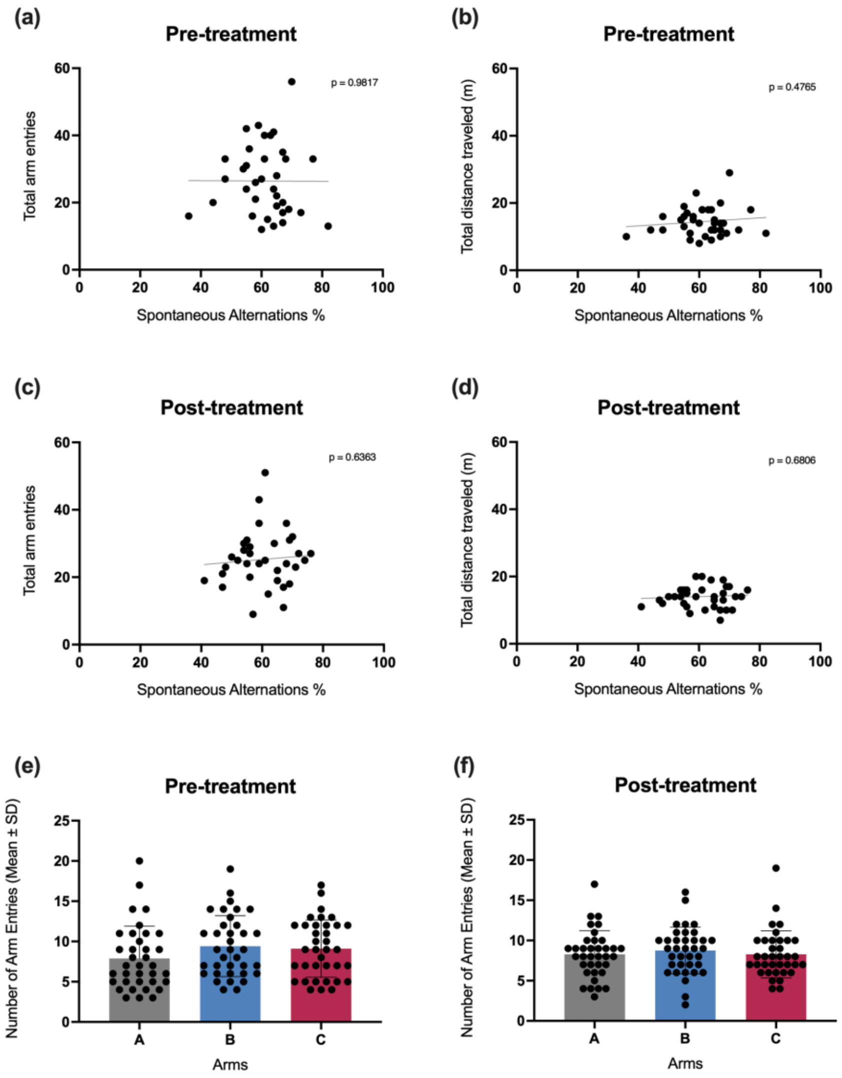


**Supplementary Figure 4. Quality control of Y-maze results.**
Pearson correlation analyses were performed across all mice (5XFAD transgenic (Tg), *n* = 21; wild-type (WT), *n* = 15; total *n* = 36) during the pre-treatment **(a, b)** and post-treatment **(c, d)** periods to examine relationships between spontaneous alternation percentage and number of arm entries **(a, c)**, and between spontaneous alternation percentage and total distance traveled **(b, d)**. No significant correlations were found, indicating that hyperdynamic locomotion did not influence the cognitive endpoint. Pearson's *r* and $R^2$ for each panel: **(a)** $r = -0.003959$, $R^2 = 1.567 \times 10^{-5}$; **(b)** $r = 0.1225$, $R^2 = 0.01501$; **(c)** $r = 0.08155$, $R^2 = 0.006651$; **(d)** $r = 0.07103$, $R^2 = 0.005046$. Total arm entries were assessed as a measure of exploratory behaviour during the pre-treatment **(e)** and post-treatment **(f)** periods. One-way ANOVA revealed no significant differences in arm entries, suggesting no environmental bias in the test.

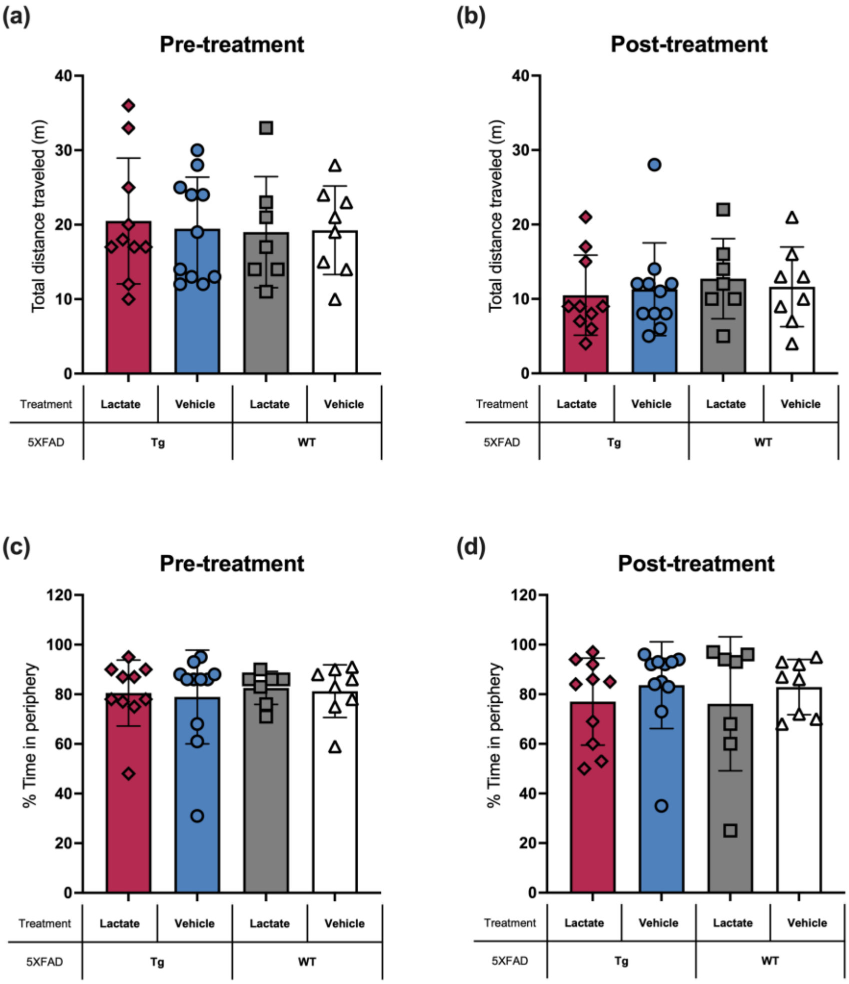


**Supplementary Figure 5. Ambulatory behaviour and thigmotaxis in the Open-field test.** Ambulation was quantified as total distance traveled (meters) during the pre-treatment **(a)** and post-treatment **(b)** periods. Thigmotaxis, assessed as the percentage of time spent in the periphery of the OFT arena, was measured during the pre-treatment **(c)** and post-treatment **(d)** periods. Experimental groups consisted of 5XFAD transgenic (Tg) mice [lactate ($n$ = 10), vehicle ($n$ = 11)] and wild-type (WT) mice [lactate ($n$ = 7), vehicle ($n$ = 8)], for a total of $n$ = 36. One-way ANOVA revealed no significant differences in locomotion or thigmotaxis across groups. Data are shown as individual data points with mean ± standard deviation (SD).

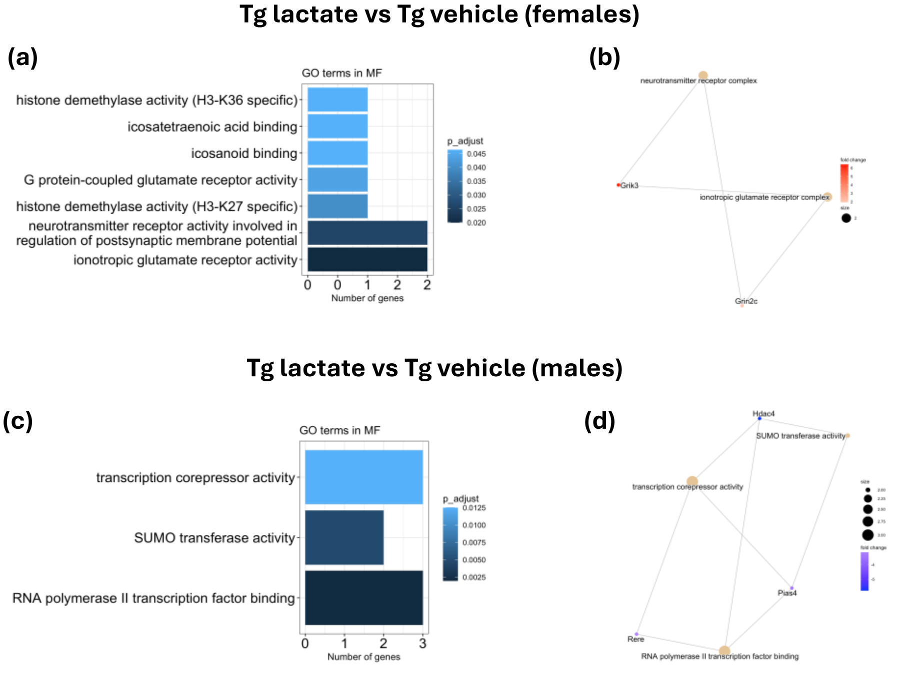


**Supplementary Figure 6. Gene Ontology (GO) enrichment analysis of differentially expressed genes (DEGs) in 5XFAD transgenic (Tg) mice following lactate treatment**

Significantly enriched GO terms within the Molecular Function (MF) category are shown for female **(a, b)** and male **(c, d)** Tg mice treated with lactate or vehicle. **(a, c)** Bar plots display significantly enriched GO terms (adjusted *p* < 0.05) with enrichment scores ($\log_2$ adjusted *p* value). **(b, d)** Network plots show relationships between enriched GO terms and associated DEGs. Node colours indicate direction of gene expression change (red = upregulated, blue = downregulated). In female Tg mice, significant enrichment was observed for GO terms related to chromatin modification and synaptic signaling, including histone demethylase activity (H3-K36 specific, H3-K27 specific), icosatetraenoic acid binding, icosanoid binding, G protein-coupled glutamate receptor activity, neurotransmitter receptor activity involved in regulation of postsynaptic membrane potential, and ionotropic glutamate receptor activity. These results suggest potential modulation of epigenetic regulation and excitatory neurotransmission by lactate in females. In male Tg mice, significant enrichment was detected for GO terms including SUMO transferase activity, transcription corepressor activity, and RNA polymerase II transcription factor binding. These functions are associated with post-translational modification, transcriptional repression, and gene regulatory control, processes implicated in neuronal stress responses and neurodegeneration.

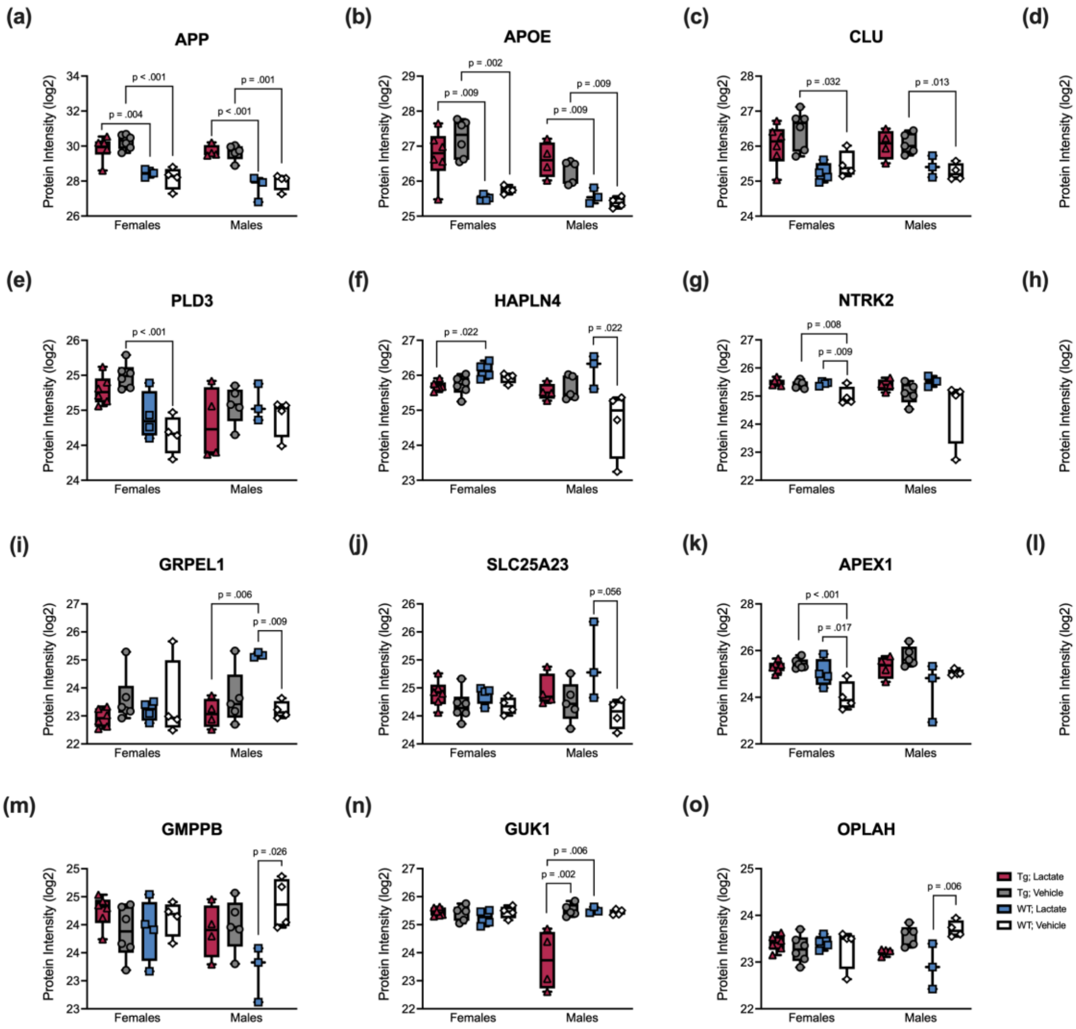


**Supplementary Figure 7. Lactate treatment alters expression of Alzheimer's disease (AD) markers, neuronal proteins, mitochondrial proteins, and cell metabolism proteins in 5XFAD transgenic (Tg) and wild-type (WT) mice.**

The experimental groups consisted of Tg mice receiving lactate (*n* = 10, 6 females and 4 males), Tg mice receiving vehicle (*n* = 11, 6 females and 5 males), WT mice receiving lactate (*n* = 7, 4 females and 3 males), and WT mice receiving vehicle (*n* = 8, 4 females and 4 males), with early and late treatment start combined. Boxplots illustrate expression levels of selected proteins across groups, including AD-related markers (APP, APOE, CLU, GFAP, PLD3), neuronal proteins (HAPLN4, NTRK2), mitochondrial proteins (ATP5G2, GRPEL1, SLC25A23), and proteins involved in cell metabolism and stress responses (APEX1, FDPS, GMPPB, GUK1, OPLAH). At the neuronal level, lactate increased HAPLN4 in WT males compared to vehicle controls ($p = 0.022$, $d = 2.63$) and NTRK2 in WT females compared to vehicle controls ($p = 0.009$, $d = 2.65$). Among mitochondrial proteins, lactate increased ATP5G2 in Tg females ($p = 0.026$, $d = 1.85$), GRPEL1 in WT males ($p = 0.009$, $d = 3.04$), and showed a nearly-significant increase of SLC25A23 in WT males ($p = 0.056$, $d = 2.21$). For metabolic and stress-related proteins, lactate significantly increased APEX1 in WT females ($p = 0.017$, $d = 2.41$), had no significant effect on FDPS, decreased GMPPB in WT males ($p = 0.026$, $d = 2.54$), decreased GUK1 in Tg males compared to Tg vehicle ($p = 0.002$, $d = 3.24$), and decreased OPLAH in WT males ($p = 0.006$, $d = 3.19$). Data are presented as box-and-whisker plots (median, interquartile range, and all individual data points). Statistical analyses were conducted using one-way ANOVA followed by Tukey's method.

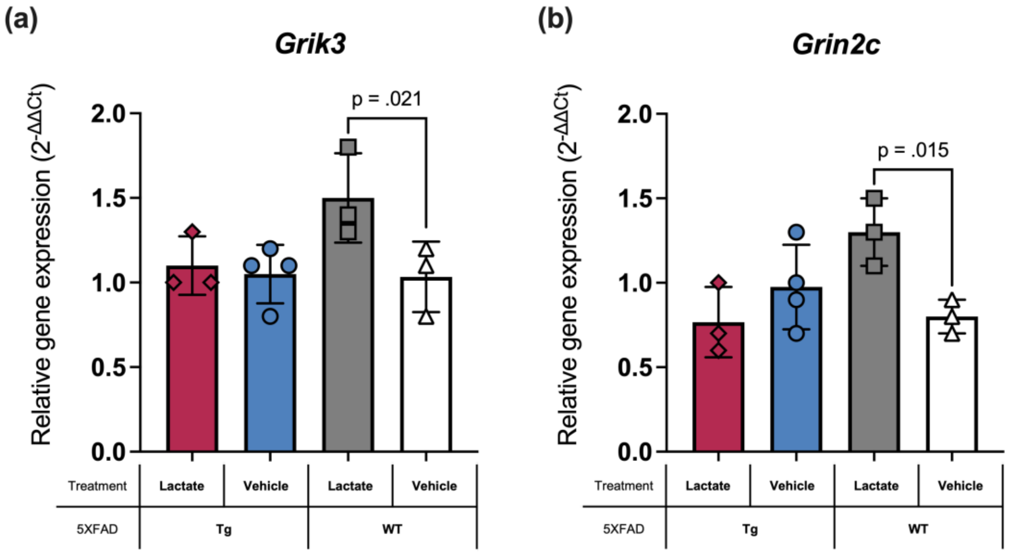


**Supplementary Figure 8. Lactate treatment upregulates *Grik3* and *Grin2c* mRNA expression in wild-type (WT) mice with late-treatment start.**

Lactate treatment significantly increased *Grik3* and *Grin2c* mRNA expression in WT mice but not 5XFAD transgenic (Tg) mice, consistent with RNA-seq findings in female mice. Relative mRNA levels were measured by RT-qPCR in Tg and WT mice receiving lactate or vehicle in the late-treatment start group. Tg lactate ($n$ = 3), Tg vehicle ($n$ = 4), WT lactate ($n$ = 3), and WT vehicle ($n$ = 3). **(a)** *Grik3* expression was significantly upregulated in WT mice treated with lactate compared to WT vehicle-treated controls ($p$ = 0.021, $d$ = 2.28). **(b)** *Grin2c* expression was significantly elevated in WT lactate-treated mice compared to WT vehicle-treated controls ($p$ = 0.015, $d$ = 2.45). Statistical analyses were performed using one-way ANOVA followed by Fisher's least significant difference test (LSD). Data are shown as individual data points with mean relative gene expression ($2^{-\Delta\Delta Ct}$) ± standard deviation (SD).

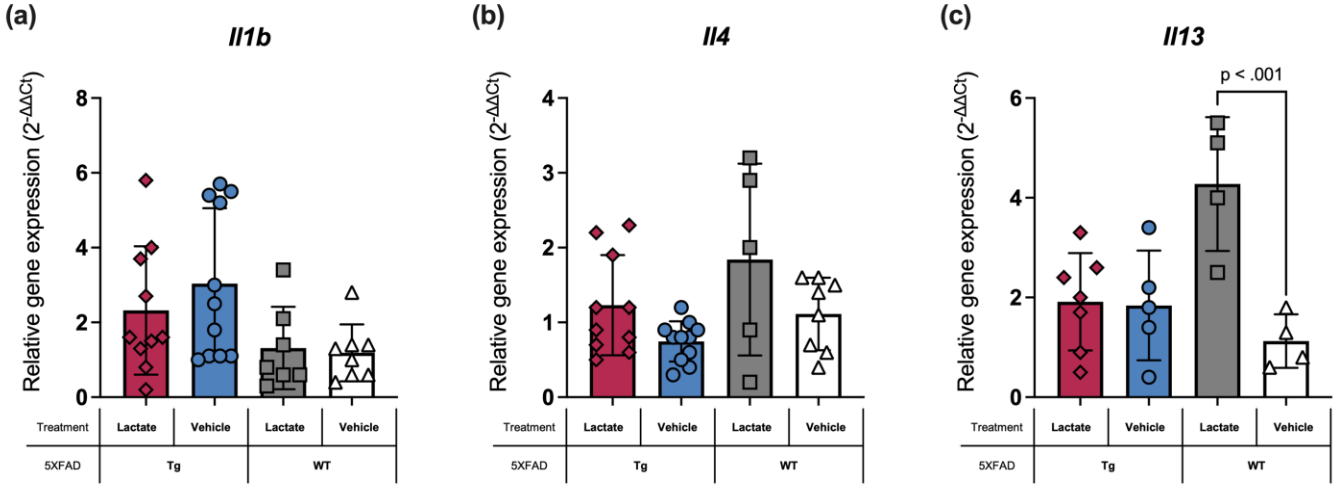


**Supplementary Figure 9. Lactate treatment increases anti-inflammatory cytokines in wild-type (WT) mice.**

Lactate treatment modulated anti-inflammatory cytokine expression in WT but not 5XFAD transgenic (Tg) mice, with a significant increase in *Il13* mRNA in WT mice, suggesting a protective effect. Relative gene expression was quantified by RT-qPCR using the $2^{-\Delta\Delta Ct}$ method. Data are presented as individual data points, with males and females combined. **(a)** No significant changes in *Il1b* mRNA expression were observed between treatment groups. **(b)** *Il4* mRNA expression exhibited an upward trend in lactate-treated WT mice compared to WT vehicle controls ($n = 5$), though the difference was not statistically significant. **(c)** *Il13* mRNA expression was significantly upregulated in lactate-treated WT mice ($n = 4$) compared to WT vehicle controls ($n = 4$, $p < 0.001$, $d = 3.07$). Statistical analyses were performed using one-way ANOVA, followed by Fisher's least significant difference test (LSD). Data are shown as individual data points with mean relative gene expression ($2^{-\Delta\Delta Ct}$) ± standard deviation (SD).

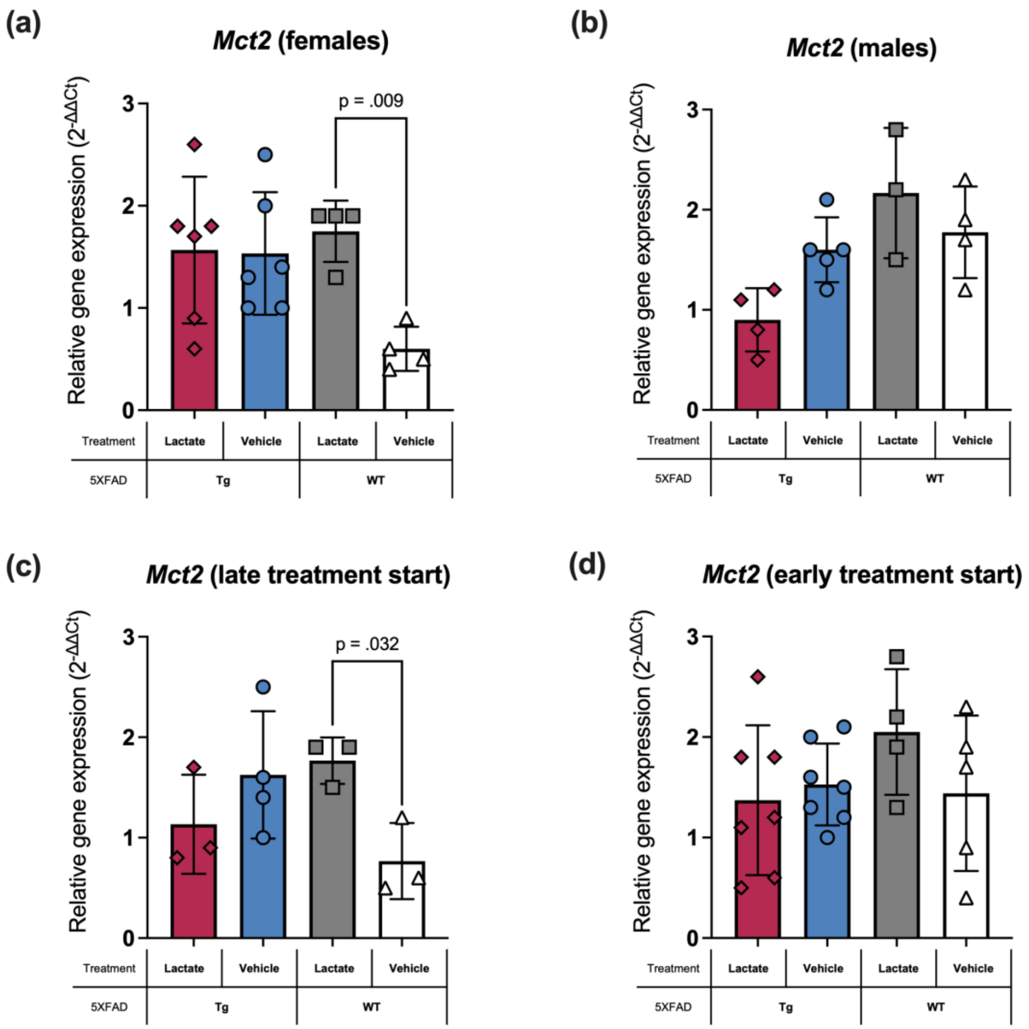


**Supplementary Figure 10. Lactate treatment increases monocarboxylate transporter 2 *(Mct2, Slc16a7)* expression in wild-type (WT) mice, specifically in females and in the late-treatment cohort.**

Relative *Mct2* mRNA expression was assessed by RT-qPCR in 5XFAD transgenic (Tg) and WT mice treated with lactate or vehicle. Panels **(a, b)** show both early- and late-treatment cohorts stratified by sex, while panels **(c, d)** show both sexes stratified by treatment start. **(a)** Lactate significantly increased *Mct2* expression in WT females compared to WT vehicle-treated controls ($n$ = 4 per group, $p$ = 0.009, $d$ = 2.11), with no significant differences in Tg mice ($n$ = 6 per group). **(b)** No significant differences were observed in male mice (Tg lactate $n$ = 4, Tg vehicle $n$ = 5, WT lactate $n$ = 3, WT vehicle $n$ = 4). **(c)** In the late-treatment cohort, WT mice treated with lactate showed significantly higher *Mct2* expression compared to WT vehicle-treated controls ($n$ = 3 per group, $p$ = 0.032, $d$ = 2.08), whereas no differences were observed in Tg mice (Tg lactate $n$ = 3, Tg vehicle $n$ = 4). **(d)** No significant differences were observed between treatment groups in the early-treatment cohort (Tg lactate $n$ = 7, Tg vehicle $n$ = 7, WT lactate $n$ = 4, WT vehicle $n$ = 5). Statistical analyses were performed using one-way ANOVA with Fisher's least significant difference test (LSD). Data are shown as individual data points with mean relative gene expression ($2^{-\Delta\Delta Ct}$) ± standard deviation (SD).

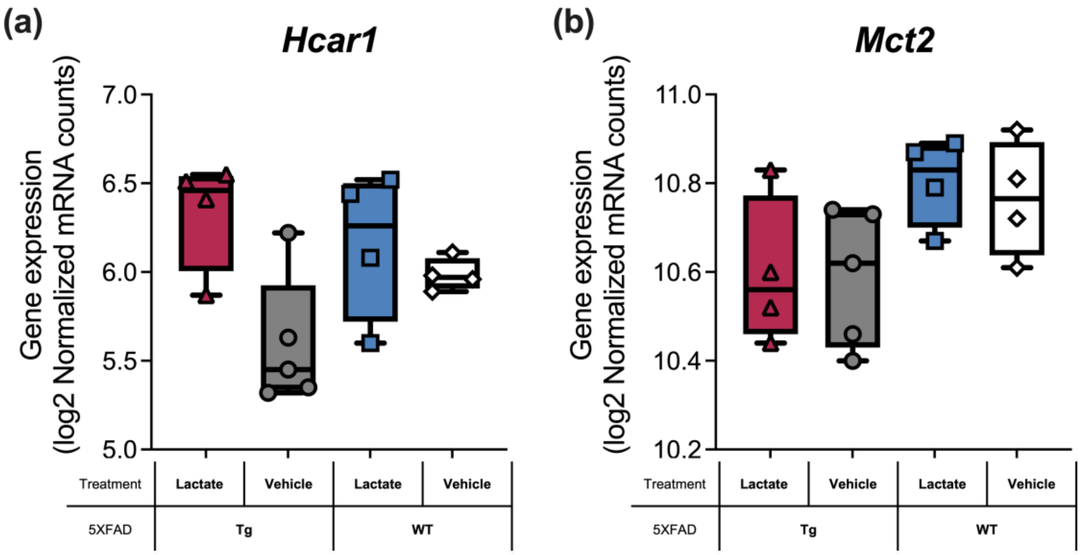


**Supplementary Figure 11. RNA-seq expression of hydroxycarboxylic acid receptor 1 (*Hcar1)* and monocarboxylate transporter 2 (*Mct2, Slc16a7).***
Transcript abundance of *Hcar1* and *Mct2* was assessed in 5XFAD transgenic (Tg) and wild-type (WT) mice treated with lactate or vehicle. Data represent the early-treatment cohort including both sexes: Tg lactate (*n* = 4), Tg vehicle (*n* = 5), WT lactate (*n* = 4), and WT vehicle (*n* = 4). DESeq2 analysis showed no significant group differences for either gene; **(a)** *Hcar1* displayed an upward trend in Tg lactate compared with Tg vehicle, whereas **(b)** *Mct2* expression was stable across conditions. However, both transcripts showed a slight trend towards lower expression in Tg than in WT on vehicle. Values are presented as box and whisker plots (median, interquartile range, and all individual data points). Statistical analysis was performed using Wald statistics with Benjamini–Hochberg correction for multiple comparisons.

**Supplementary Table 1. Primer sequences and assay characteristics for 5XFAD transgene detection of amyloid precursor protein (*App)* and presenilin-1 (*Psen1)*.** The table includes forward (FW) and reverse (RV) primers, expected amplicon sizes, melting temperatures ($T_m$), and annealing temperatures ($T_a$).

| Gene | Target | Amplicon size (bp) | Primer | DNA sequence | Melting temperature, $T_m$, (°C) | Annealing temperature, $T_a$, (°C) |
|---|---|---|---|---|---|---|
| *APP* | Tg | ~377 | oIMR3610_Jax (FW) | 5'-AGGACTGACCACTCGACCAG-3' | 86.0 ± 1.0 (74.4) | 60.5 |
| | Tg | ~377 | oIMR3611_Jax (RV) | 5'-CGGGGGTCTAGTTCTGCAT-3' | 86.0 ± 1.0 (73.8) | 60.5 |
| | Internal positive control | ~324 | oIMR7338_Jax (FW) | 5'-CTAGGCCACAGAATTGAAAGATCT-3' | 81.0 ± 1.0 (75.6) | 60.0 |
| | Internal positive control | ~324 | oIMR7339_Jax (RV) | 5'-GTAGGTGGAAATTCTAGCATCATCC-3' | 81.0 ± 1.0 (75.8) | 60.0 |
| *PSEN1* | Tg | ~608 | oIMR1644_Jax (FW) | 5'-AATAGAGAACGGCAGGAGCA-3' | 86.0 ± 1.0 (70.8) | 58.5 |
| | Tg | ~608 | oIMR1645_Jax (RV) | 5'-GCCATGAGGGCACTAATCAT-3' | 86.0 ± 1.0 (74.4) | 58.5 |
| | Internal positive control | ~324 | oIMR7338_Jax (FW) | 5'-CTAGGCCACAGAATTGAAAGATCT-3' | 81.0 ± 1.0 (75.6) | 53.5 |
| | Internal positive control | ~324 | oIMR7339_Jax (RV) | 5'-GTAGGTGGAAATTCTAGCATCATCC-3' | 81.0 ± 1.0 (75.8) | 53.5 |
